# Molecular basis and design principles of a system for switchable front-rear polarity and directional migration

**DOI:** 10.1101/2022.12.09.519731

**Authors:** Luís António Menezes Carreira, Dobromir Szadkowski, Stefano Lometto, Georg K.A. Hochberg, Lotte Søgaard-Andersen

## Abstract

During cell migration, front-rear polarity is spatiotemporally regulated; however, the underlying design of regulatory interactions vary. In rod-shaped *Myxococcus xanthus* cells, a spatial toggle switch dynamically regulates front-rear polarity. The polarity module establishes front-rear polarity by guaranteeing front pole-localization of the small GTPase MglA. Conversely, the Frz chemosensory system, by acting on the polarity module, causes polarity inversions. MglA localization depends on the RomR/RomX GEF and MglB/RomY GAP complexes that localize asymmetrically to the poles by unknown mechanisms. Here, we show that RomR and the MglB and MglC roadblock domain proteins generate a positive feedback by forming a RomR/MglC/MglB complex, thereby establishing the rear pole with high GAP activity that is non-permissive to MglA. MglA at the front engages in negative feedback that inhibits the RomR/MglC/MglB positive feedback allosterically, thus ensuring low GAP activity at this pole. These findings unravel the design principles of a system for switchable front-rear polarity.

## Introduction

Cell polarity with the asymmetric localization of proteins within cellular space is ubiquitous and foundational for many cellular functions, including growth and motility^1-3^. Nevertheless, how polarity emerges at cellular scales from local protein-protein interactions and how it is dynamically controlled is poorly understood. Polarity regulators are often connected to generate networks that include positive feedback, negative feedback and/or mutual inhibition^2, 4-7^. In transcriptional regulation, it is well-established that different designs of regulatory circuits can result in functionally equivalent outcomes, e.g. double-negative is functionally equivalent to double-positive regulation^8^. Similarly, polarity-regulating networks with functionally equivalent outcomes can have different designs, raising the question of why a particular network design has been selected.

A recurring theme in polarity-regulating systems is the localization of the active GTP-bound form of a small GTPase at a single intracellular location^6, 7, 9-12^. The GTPase, in turn, interacts with downstream effectors to implement a specific response. These GTPases are molecular switches that alternate between an inactive, GDP-bound and an active, GTP-bound conformation^13^. The activation/deactivation cycle is regulated by a cognate guanine-nucleotide exchange factor (GEF), which facilitates the exchange of GDP for GTP, and a GTPase activating protein (GAP), which stimulates the low intrinsic GTPase activity^14^. Two experimentally and theoretically well-studied systems illustrate how polarity-regulating networks with different designs can result in equivalent outcomes. In *Saccharomyces cerevisiae* lacking the small GTPase Rsr1, the location of the single bud site depends on where the GTPase Cdc42 spontaneously forms a single cluster on the membrane. The responsible regulatory network centers on at least one positive feedback directly involving Cdc42^4, 9^. Briefly, Cdc42-GTP spontaneously forms a cluster on the membrane and then recruits a complex that includes the GEF Cdc24^9^. Because Cdc24 activates additional Cdc42, Cdc24 recruitment stimulates the accumulation of additional Cdc42-GTP, closing the positive feedback^9^. Cdc42 GAPs inhibit Cdc42 cluster growth and may be part of a negative feedback^9, 15, 16^. In the alternative system, unidirectional migration of the rod-shaped cells of the bacterium *Myxococcus xanthus* depends on the localization of the GTPase MglA at the leading front pole. In this case, the positive feedback does not involve MglA but rather the GAP MglB and the RomR scaffold^17^. Ultimately, these two proteins establish a rear, lagging pole with high GAP activity leaving only the opposite pole free to recruit MglA-GTP^17^. Thus, both systems generate a single Cdc42/MglA cluster. Here, we focus on the mechanistic basis of polarity establishment in *M. xanthus* and the functional properties conferred by the underlying network compared to the circuit that brings about Cdc42 cluster formation.

*M. xanthus* migrates unidirectionally on surfaces using two motility machines that assemble at the leading pole^11, 18, 19^. In response to signalling by the Frz chemosensory system, cells reverse the direction of movement^20^. During reversals, cells invert polarity and the pole at which the motility machines assemble switches^21, 22^. Motility and its regulation by the Frz system are essential for multicellular morphogenesis with the formation of predatory colonies and spore-filled fruiting bodies^11, 18, 19^. Active MglA-GTP stimulates the assembly of the motility machineries at the leading cell pole^23-25^. Front-rear polarity is regulated dynamically by two interconnected protein modules, i.e. the polarity module and the Frz chemosensory system, that in combination generate a spatial toggle switch. The polarity module sets up the leading/lagging polarity axis and, in addition to MglA, comprises four proteins that also localize asymmetrically to the cell poles (Fig. 1A). The homodimeric roadblock domain protein MglB alone has GAP activity and together with its low-affinity co-GAP RomY, forms the MglB/RomY complex, which is the active GAP *in vivo*^26-28^. RomX alone has GEF activity and forms the RomR/RomX complex, the active GEF *in vivo* that also serves as a polar recruitment factor for MglA-GTP^29^.

**Fig 1.**
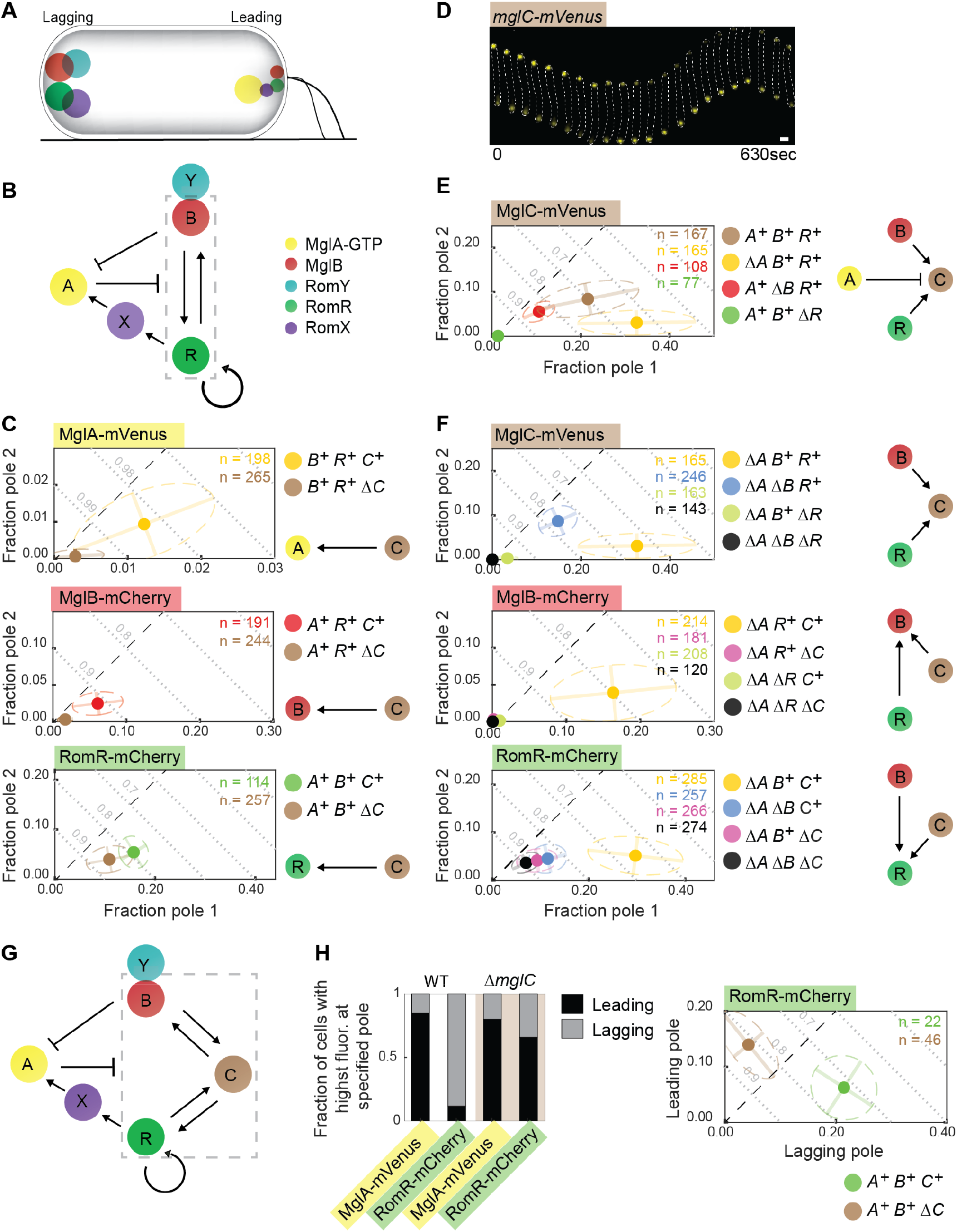
MglC localization depends on MglA, MglB and RomR, and *vice versa*. A. Schematic of MglA-GTP, MglB, RomR, RomX and RomY localization. T4P are shown at the leading pole. The size of a circle indicates the relative amount of a protein at a pole. Colour code as in Fig. 1B. B. Schematic of interactions between polarity proteins. The dashed grey box indicates the RomR/MglB positive feedback. C. MglA, MglB and RomR polar localization depends on MglC. All fusion proteins were synthesized from their native locus. In the diagrams, the poles with the highest and lowest polar fraction of fluorescence are defined as pole 1 and pole 2, respectively. The mean fraction of fluorescence at each pole is indicated by the filled circles. Dispersion of the single-cell measurements is represented by error bars and ellipses (colored dashed lines). The direction and length of error bars are defined by the eigenvectors and square root of the corresponding eigenvalues of the polar fraction covariance matrix for each strain. Black dashed lines are symmetry lines, grey dashed lines are guidelines to indicate the fraction of total polar fluorescence. Number of cells analyzed (n) is indicated in the top right corners. *mglA, mglB, mglC* and *romR* genotypes are indicated with *A, B, C* and *R*, respectively. Schematics of the effects observed is shown on the right. D. MglC-mVenus localizes asymmetrically and dynamically to the cell poles. The fusion was synthesized from the native locus. Cells were imaged by time-lapse microscopy at 30sec intervals. Scale bar, 1μm. E. MglC polar localization depends partially on MglB and strongly on RomR. Data are presented as in Fig. 1C. F. Quantification of the polar localization of MglC-mVenus, MglB-mCherry and RomR-mCherry in the absence of MglA. Data are presented as in Fig. 1C. G. MglC is a component of the RomR/MglC/MglB positive feedback. Dashed grey box, the RomR/MglC/MglB positive feedback. H. MglC is essential for establishing correct RomR polarity. Cells were imaged by time-lapse microscopy as in Fig. 1D, and the fractions of cells with the brightest cluster at the leading or lagging pole determined. Left panel, a summary of fractions of cells with indicated localization pattern. Right panel, quantification of RomR-mCherry localization in moving cells.

Experiments and mathematical modelling have uncovered an intricate set of regulatory interactions between the proteins of the polarity module^17, 26-32^ (Fig. 1B). The RomR scaffold is at the base of all other polarity proteins’ polar localization and also reinforces its own polar localization, thereby establishing a positive feedback^17^. RomR also engages in a positive feedback with MglB by an unknown mechanism^17^. Additionally, RomR directly recruits RomX to form the RomR/RomX GEF complex. High concentrations of polar MglB stimulate polar recruitment of its low-affinity interaction partner RomY^28^. At the RomR node of the RomR/MglB positive feedback, RomR/RomX promotes MglA-GTP polar recruitment (Fig. 1B – connector from RomR/RomX to MglA)^29^, and at the MglB node, MglB/RomY inhibits MglA-GTP polar recruitment (Fig. 1B – connector from MglB/RomY to MglA)^26-28^. Finally, MglA-GTP disrupts the RomR/MglB positive feedback by an unknown mechanism (Fig. 1B – connector from MglA to dashed box)^17^. Together these interactions have been suggested to result in the system’s emergent properties (Fig. 1AB)^17, 28^. Briefly, at the pole with the highest RomR concentration, the RomR/MglB positive feedback establishes a pole with high concentrations of RomR/RomX and MglB/RomY. Due to the presence of the MglB/RomY complex, GAP activity dominates over GEF activity at this pole, thus inhibiting MglA-GTP recruitment, and this pole becomes the lagging pole. At the opposite pole, RomR/RomX GEF activity dominates over GAP activity because the low concentration of MglB is insufficient to recruit RomY^28^. Consequently, MglA-GTP is recruited to this pole and engages in the negative feedback to inhibit the RomR/MglB positive feedback, thereby maintaining the low concentration of the other polarity regulators. The Frz system is the second module of the spatial toggle switch, and the polarity module is the downstream target of this system. Frz signaling causes the inversion of polarity of the proteins of the polarity module by an unknown mechanism, thus laying the foundation for assembly of the motility machineries at the new leading pole^30, 31, 33, 34^.

Among the interactions of the proteins of the polarity module, the positive feedback of RomR on itself, the RomR/MglB positive feedback, and the inhibitory effect of MglA-GTP on this positive feedback are poorly understood. MglC is also a homodimeric roadblock domain protein^35-37^ and is involved in cell polarity regulation by an unknown mechanism^36^. Because MglC interacts with RomR and MglB^35, 36^, MglC was a candidate for acting in the RomR/MglB positive feedback. Here, we show that MglC forms a complex with RomR and MglB, thereby establishing a RomR/MglC/MglB positive feedback and that MglA-GTP inhibits this positive feedback by breaking the interaction between the MglC and MglB roadblock domain proteins. Moreover, we demonstrate that the RomR/MglC/MglB positive feedback lays the foundation for switchable polarity.

## Results

### MglC is important for Frz-induced cellular reversals

To investigate the function of MglC in polarity, we recharacterized the motility defects of a mutant with an in-frame deletion of *mglC* (Δ*mglC*). In agreement with previous findings^36^, the Δ*mglC* mutant has defects in both gliding and T4P-dependent motility in population-based motility assays, and ectopic expression of *mglC* complemented these defects (Fig. S1AB). In single cell-based motility assays (Fig. S1C), and consistent with previous observations^36^, Δ*mglC* cells moved with the same speed as wild-type (WT) for both motility systems; however, similarly to the Δ*frzE* negative control that lacks the FrzE kinase, Δ*mglC* cells had a significantly lower reversal frequency than WT.

To discriminate whether the Δ*mglC* mutant is unresponsive to or has reduced sensitivity to Frz signaling, we treated WT and Δ*mglC* cells with the short-chain alcohol isoamyl alcohol (IAA) that highly stimulates reversals in a FrzE-dependent manner^38^. WT and the Δ*mglC* mutant responded similarly to 0.3% IAA with the formation of colonies that had smooth edges and no visible flares on 0.5% agar, which is optimal for T4P-dependent motility, and few single cells at the edge on 1.5% agar, which is optimal for gliding motility (Fig. S1A). Such smooth colony edges indicate a high reversal frequency^20, 39^. We conclude that the Δ*mglC* mutant does not have a defect in motility *per se* but reduced sensitivity to Frz signaling resulting in a reduced reversal frequency.

### MglC is important for the polar localization of MglA, MglB and RomR

Because the polarity module is the downstream target of the Frz system, we quantified the polar localization of active, fluorescently labelled fusions of the polarity proteins in the absence of MglC. Because RomX localization follows that of RomR^29^ and RomY localization follows the highest concentration of MglB^28^, we used the RomR and MglB localization as readouts for the localization of the RomR/RomX complex and MglB/RomY complex, respectively.

In snapshots of Δ*mglC* cells (Fig. 1C), polar localization of MglA-mVenus and MglB-mCherry was strongly reduced, while RomR-mCherry polar localization was only partially lost. MglA, MglB and RomR accumulated independently of MglC (Fig. S1D).

### MglC polar localization depends partially on MglB and strongly on RomR

To study MglC localization, we first observed that a fully active MglC-mVenus fusion expressed from the native site (Fig. S1AB) localized in a bipolar asymmetric pattern with a large cluster at the lagging pole in WT cells and switched polarity during reversals (Fig. 1D). The bipolar asymmetric pattern was also evident in snapshots (Fig. 1E). In the absence of MglA, MglC-mVenus was more polar (Fig. 1E). However, in the absence of MglB, MglC-mVenus polar localization was partially lost; and, in the absence of RomR, it was almost completely lost (Fig. 1E). MglC-mVenus accumulated independently of MglA, MglB and RomR (Fig. S1E). Thus, MglA inhibits MglC polar localization while MglC depends partially on MglB and strongly on RomR. Of note, in the absence of RomR, MglB fails to support significant MglC polar localization.

### MglC establishes the RomR/MglC/MglB positive feedback

Because the interpretation of the results for polar localization of MglC, MglB and RomR can be challenging due to the inhibitory effect of MglA-GTP on the RomR/MglB positive feedback, we quantified their polar fluorescence in strains lacking MglA.

MglC-mVenus polar localization in the Δ*mglA*Δ*mglB* mutant was partially lost compared to the Δ*mglA* mutant, almost completely abolished in the Δ*mglA*Δ*romR* mutant, and completely abolished in the triple Δ*mglA*Δ*mglB*Δ*romR* mutant (Fig. 1F). These observations confirm that MglC-mVenus polar localization depends partially on MglB and strongly on RomR. They also confirm that in the absence of RomR, MglB fails to support MglC polar localization significantly.

MglB-mCherry polar localization in the Δ*mglA*Δ*mglC*, the Δ*mglA*Δ*romR* and the Δ*mglA*Δ*mglC*Δ*romR* mutants was almost completely lost (Fig. 1F). These results confirm that MglB-mCherry polar localization depends strongly on MglC and, as previously shown^17^, on RomR. Moreover, neither MglC nor RomR alone can establish efficient polar MglB-mCherry localization.

RomR-mCherry polar localization was partially abolished in the Δ*mglA*Δ*mglB*, Δ*mglA*Δ*mglC* and Δ*mglA*Δ*mglB*Δ*mglC* mutants (Fig. 1F). Thus, both MglB and MglC are important but not essential for RomR polar localization. Moreover, neither MglB nor MglC alone further stimulates RomR polar localization.

These observations demonstrate that RomR alone localizes polarly, and they support that RomR recruits MglC, which then recruits MglB. The observations that (1) MglB stimulates MglC polar localization in the presence of RomR, and (2) MglB together with MglC stimulates RomR polar localization support that the three proteins establish a positive feedback that reinforces their polar localization (Fig. 1G). These observations also suggest that the previously established RomR/MglB positive feedback depends on MglC, i.e. MglC helps to generate a RomR/MglC/MglB positive feedback by acting between RomR and MglB (Fig. 1G). Because MglA inhibits the RomR/MglB positive feedback^17^, this model also explains the observation that MglA inhibits MglC polar localization (Fig. 1E). Moreover, the reduced MglA polar localization in the absence of MglC (Fig. 1C) is a direct outcome of the reduced RomR polar localization in the absence of MglC.

To further test the idea of the RomR/MglC/MglB positive feedback, we leveraged an established approach to monitor the cooperative polar recruitment of RomR-mCherry^17^. In this approach, a vanillate-inducible promoter drives *romR-mCherry* expression; upon induction, RomR-mCherry polar localization is followed by time-lapse fluorescence microscopy. To monitor RomR-mCherry synthesis over time, we estimate the RomR-mCherry concentration in individual cells, referred to as the fluorescence concentration, by measuring total cellular fluorescence and then normalizing by cell area, which we use as a proxy for cell volume.

Upon induction of *romR-mCherry* expression in the Δ*mglA*Δ*mglB*Δ*romR*Δ*mglC* quadruple mutant (Fig. S2A) and the Δ*mglA*Δ*romR*Δ*mglC* triple mutant (Fig. S2B), RomR-mCherry localized asymmetrically to the poles at all fluorescence concentrations and quantitatively followed the pattern previously observed in the Δ*mglA*Δ*mglB*Δ*romR* triple mutant (Fig. S2C). As described^17^, the observations that the fractions of RomR-mCherry at both poles increase with fluorescence concentration at low induction levels provide evidence for positive cooperativity in RomR-mCherry polar localization. Because RomR-mCherry polar localization is quantitatively similar in these three strains, we conclude that MglC, similar to MglB, is not essential for the positive feedback of RomR on itself. By contrast, in the Δ*mglA*Δ*romR* double mutant, RomR-mCherry polar localization was increased and more asymmetric, with the brighter pole accounting for a larger fraction of RomR-mCherry fluorescence (Fig. S2D). This observation confirms that MglC is essential for establishing the RomR/MglB positive feedback and that the RomR/MglB positive feedback is, in fact, a RomR/MglC/MglB positive feedback (Fig. 1G).

### MglC is essential for establishing correct RomR polarity

The model for polarity establishment (Fig. 1G) predicts that in the absence of MglC and, therefore, the RomR/MglC/MglB positive feedback, the residual polar RomR, together with RomX, will recruit MglA-GTP. As a result, MglA-GTP and RomR/RomX will have their highest polar fluorescence at the leading pole. To test this prediction, we performed time-lapse fluorescence microscopy of moving cells. In WT, MglA-mVenus localized with a large cluster at the leading pole and RomR-mCherry with a large cluster at the lagging pole in most cells (Fig. 1H). Importantly, and as predicted, in the Δ*mglC* mutant, MglA-mVenus and RomR-mCherry had their highest polar fluorescence at the leading pole in most cells (Fig. 1H). We conclude that MglC is important not only for the polar localization of MglA, MglB and RomR but also for establishing the correct polarity of RomR-mCherry.

### RomR, MglC and MglB interact to form a complex

To investigate the mechanism underlying the RomR/MglC/MglB positive feedback, we tested for direct interactions between RomR, MglC, MglB and MglA using pull-down experiments *in vitro* with purified proteins. In agreement with previous observations in *in vitro* pull-down experiments^35^ and Bacterial Adenylate Cyclase-Based Two-Hybrid (BACTH) assays^36^, Strep-MglC pulled-down His_6_-MglB and MalE-RomR in pairwise combinations, but not MglA-His_6_ preloaded with GTP (Fig. 2A; Fig. S3A). In pairwise combinations using MalE-RomR as a bait, MalE-RomR pulled-down Strep-MglC but not His_6_-MglB; notably, in the presence of all three proteins, RomR-MalE pulled down Strep-MglC as well as His_6_-MglB (Fig. 2B; Fig. S3B). Finally, in pairwise combinations, His_6_-MglB pulled-down Strep-MglC but not MalE-RomR; however, in the presence of all three proteins, His_6_-MglB pulled-down Strep-MglC as well as MalE-RomR (Fig. 2C; Fig S3C).

**Fig. 2.**
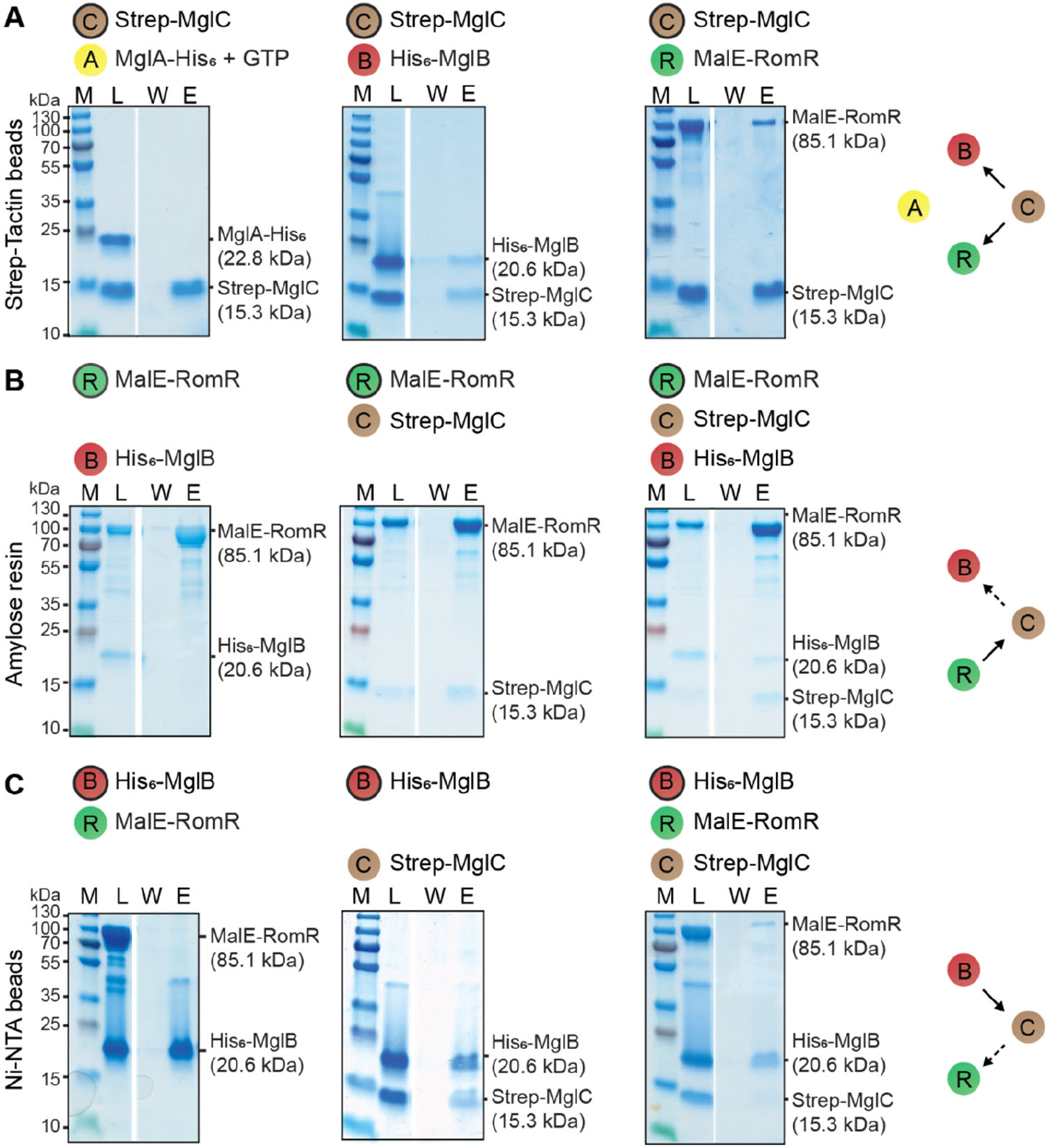
MglB, MglC and RomR form a complex *in vitro*. A-C. Proteins were mixed at final concentrations of 10µM and applied to the indicated matrices. Matrices were washed and bound proteins eluted. The bait protein is indicated by the black circle. In experiments with MglA-His_6_, the protein was preloaded with GTP and all buffers contained 40µM GTP. Equivalent volumes of the load (L), last wash (W) and elutate (E) were separated on the same SDS-PAGE gel and stained with Coomassie Brilliant Blue. Gap between lanes indicates lanes deleted for presentation purposes. Calculated molecular masses of the indicated proteins are indicated on the right and molecular weight markers (M) on the left. In the schematics on the right, a direct interaction with the bait is indicated by a black and an indirect interaction by a dashed arrow.

Next, we determined whether MglC and/or RomR/MglC have MglA GAP activity or interfere with MglB and/or MglB/RomY GAP activity. To this end, we determined MglA-His_6_ GTPase activity in the presence of RomR, MglC, MglB and/or RomY. Neither Strep-MglC nor RomR/MglC affected MglA GTPase activity in the presence or absence of MglB-His_6_ and/or Strep-RomY (Fig. S4).

We conclude that MalE-RomR, Strep-MglC and His_6_-MglB interact to form a complex in which Strep-MglC is sandwiched between MalE-RomR and His_6_-MglB.

### The MglB KRK surface region represents the interface for interaction with MglC

To elucidate the structural basis for the RomR→MglC→MglB interactions, we took advantage of structural information for MglA, MglB and MglC^35, 40-42^. Each MglB protomer in the homodimer consists of a five-stranded β-sheet sandwiched between the α2-helix and the α1/α3-helices. In the dimer, the α2-helices generate the so-called two-helix side and the pairs of α1/α3-helices the so-called four-helix side (Fig. 3A). In the crystallographic structure of the MglA-GTPɣS:MglB_2_ complex, the MglA monomer interacts asymmetrically with the two-helix side of the MglB dimer^40-42^.

**Fig. 3.**
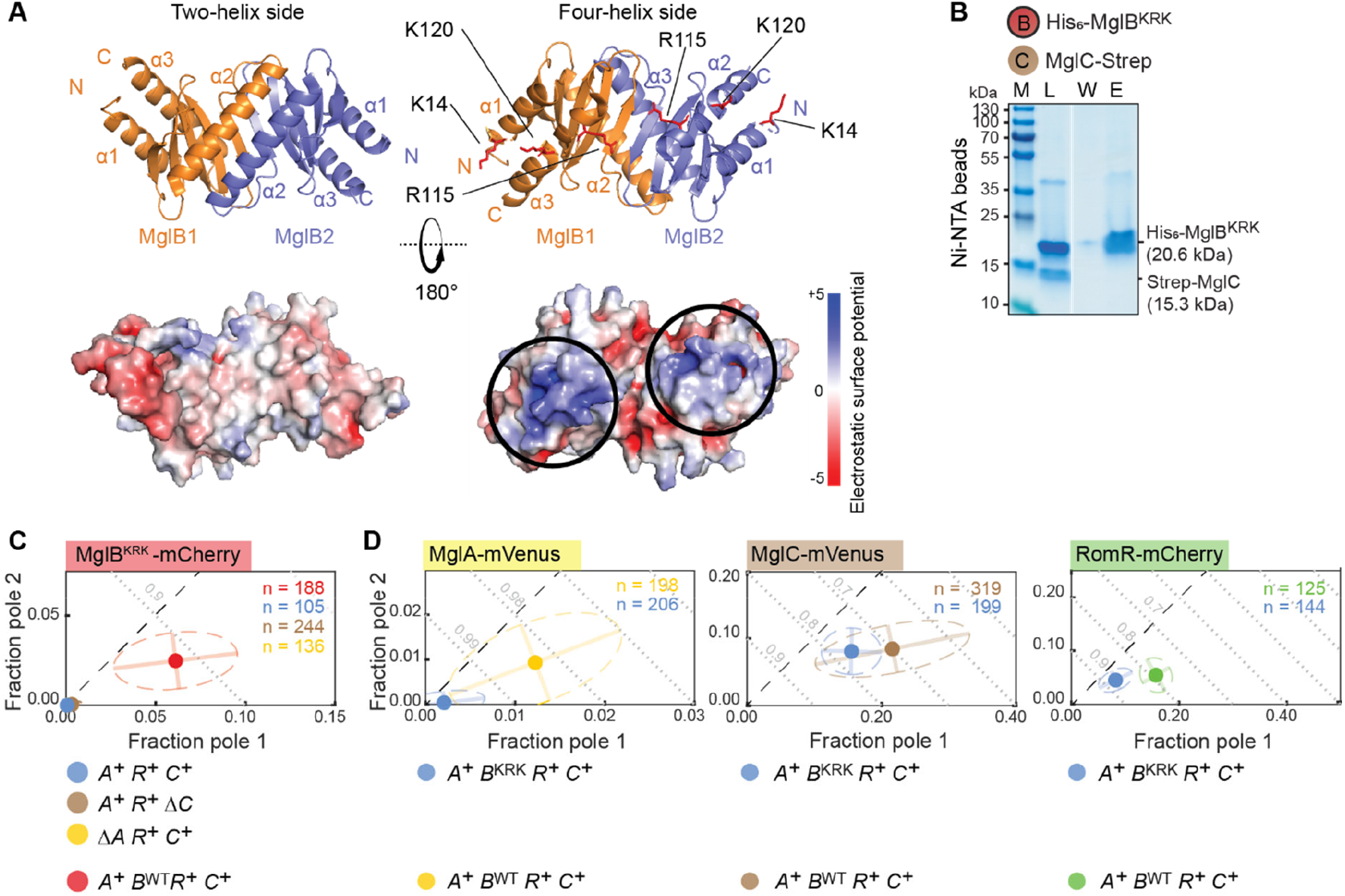
The MglB KRK surface regions represent the interface for interaction with MglC. A. Crystallographic structure of MglB dimer (pdb ID: 6hjm^40^) viewed from the two-helix and four-helix sides. Lower panels, surface representation of MglB dimer based on electrostatic surface potential contoured from +5 to -5 kT e^-1^. The K14, R115 and K120 residues are indicated in red on the four-helix side and the corresponding positively charged surface regions by black circles in the electrostatic surface potential diagrams. B. The MglB^KRK^ variant does not interact with MglC. Pull-down experiment was performed with His_6_-MglB^KRK^ as bait on the indicated resin and the data presented as in Fig. 2. C. MglB^KRK^-mCherry has reduced polar localization. For comparison, MglB^WT^-mCherry is included (red dot). MglB^KRK^-mCherry was synthesized from the native locus. D. MglB^KRK^ causes reduced polar localization of MglA, MglC and RomR. For comparison, the localization of the three fusion proteins are included in the presence of MglB^WT^ (yellow, brown and green dots). MglB^KRK^ was synthesized from the native locus. In C-D, data are presented as in Fig 1C.

Based on conservation, Galicia *et al*. reported the MglB^K14A R115A K120A^ variant (henceforth MglB^KRK^) with three substitutions in two positively charged surface regions on the four-helix side of the dimer (Fig. 3A). This variant has GAP activity but localizes diffusely by an unknown mechanism^40^. We hypothesized that the positively charged surface regions in the MglB dimer defined by the K14, R115, K120 residues (Fig. 3A) could be involved in the interaction between MglB and MglC.

In *in vitro* pull-down experiments, His_6_-MglB^KRK^ did not detectably bind Strep-MglC (Fig. 3B). Consistently, polar localization of MglB^KRK^-mCherry in otherwise WT cells was strongly reduced independently of the presence or absence of MglC and MglA (Fig. 3C; Fig. S5A). In the inverse experiment, MglB^KRK^ caused a strong reduction in MglA-mVenus localization, while MglC-mVenus and RomR-mCherry polar localization was partially abolished (Fig. 3D; Fig. S5A). We conclude that MglB^KRK^ is deficient in interacting with MglC and infer that the positively charged KRK surface regions in the MglB dimer represent the interface to MglC.

### The MglC FDI surface region represents the interface for interaction with MglB

The MglC homodimer’s structure is similar to that of MglB with two-helix and four-helix sides (Fig. 4A)^35^. Based on conservation, McLoon *et al*. reported the MglC^F25A D26A I28A^ variant (henceforth MglC^FDI^) with substitutions on the two-helix side (Fig. 4A; Fig. S6). In the dimer, the regions defined by the F25, D26, I28 residues are separated and describe two negatively charged surface regions (Fig. 4A). The FDI substitutions were reported to weaken the MglB/MglC interaction but not the MglC/RomR interaction based on BACTH assays^36^ supporting that the two FDI surface regions define the interaction interface of MglC to MglB.

**Fig. 4.**
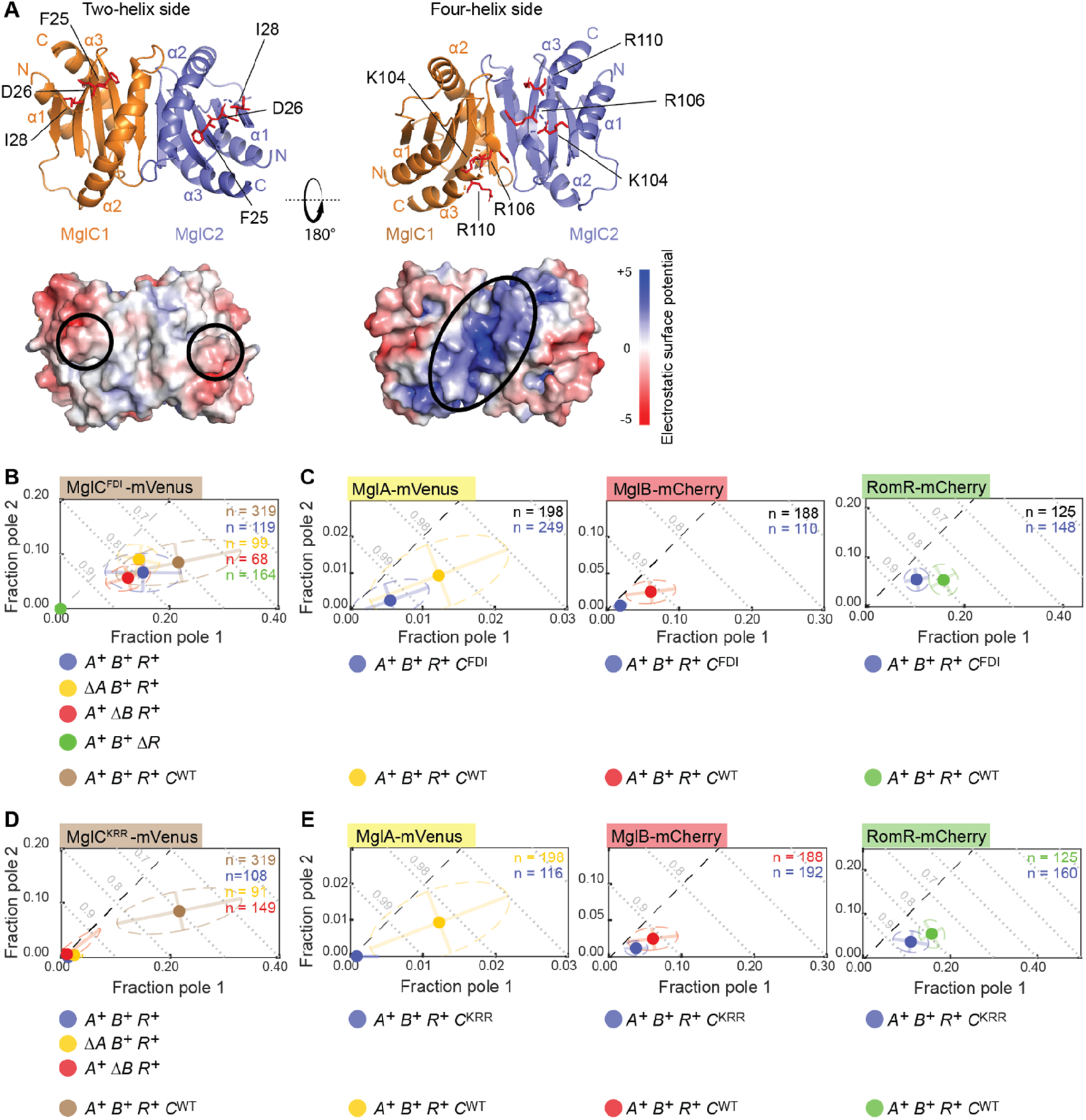
The MglC FDI and KRR surface regions represent the interfaces for interaction with MglB and RomR, respectively. A. Crystallographic structure of MglC dimer (pdb ID: 7ct3^35^) viewed from two-helix and four-helix sides. Lower panels, surface representation of MglC dimer based on electrostatic surface potential contoured from +5 to -5 kT e^-1^. The F25, D26 and I28 residues and the K104, R106, R110 residues are indicated in red on the two-helix and the four-helix sides, respectively, and the corresponding negatively and positively charged surface regions indicated by black circles in the electrostatic surface potential diagrams. B. MglC^FDI^-mVenus has reduced polar localization. For comparison, MglC^WT^-mVenus is included (brown dot). MglC^FDI^-mVenus was synthesized from the native locus. C. MglC^FDI^ causes reduces polar localization of MglA, MglB and RomR. For comparison, the localization of the three fusion proteins is included in the presence of MglC^WT^ (yellow, red and green dots). MglC^FDI^ was synthesized ectopically. D. MglC^KRR^-mVenus has strongly reduced polar localization. For comparison, MglC^WT^-mVenus is included (brown dot). MglC^KRR^-mVenus was synthesized from the native locus. A. MglC^KRR^ causes reduced polar localization of MglA, MglB and RomR. For comparison, the localization of the three fusion proteins is included in the presence of MglC^WT^ (yellow, red and green dots). MglC^KRR^ was synthesized ectopically. In B-E, data are presented as in Fig 1C.

We sought to verify the effect of the MglC^FDI^ variant on the MglC/MglB interaction *in vitro* but were unable to purify a soluble Strep-tagged variant. Importantly, polar localization of MglC^FDI^-mVenus in otherwise WT cells was partially lost in comparison to MglC-mVenus (Fig. 4B; Fig. S5B). Moreover, MglC^FDI^–mVenus polar localization did not change much upon removal of MglA or MglB but was abolished by removal of RomR (Fig. 4B). In the inverse experiment, MglC^FDI^, similar to the Δ*mglC* mutation, caused strong reductions in MglA-mVenus and MglB-mCherry polar localization while RomR-mCherry polar localization was partially abolished (Fig. 4C; Fig. S5B). We conclude that MglC^FDI^ is deficient in interacting with MglB but not with RomR and infer that the negatively charged FDI surface regions in the MglC dimer represent the interface to MglB.

### The MglC KRR surface region represents the interphase for interaction with RomR

In addition to the FDI residues, the K104, R106, R110 residues (MglC numbering) are highly conserved in MglC homologs (Fig. S6). These three residues are located on the four-helix side and define a continuous, positively charged, surface-exposed region in the dimer (Fig. 4A). Because this region is apart from the FDI region and MglC interacts with MglB and RomR in parallel, we hypothesized that it could interface with RomR.

To this end, we generated MglC^K104A R106A R110A^ variants (henceforth, MglC^KRR^). We were unable to purify a soluble Strep-tagged MglC^KRR^ variant. Importantly, polar localization of MglC^KRR^-mVenus in otherwise WT cells was strongly reduced compared to MglC-mVenus (Fig. 4D; Fig. S5C). In the inverse experiment, MglC^KRR^, similar to the Δ*mglC* mutation and MglC^FDI^, caused a strong reduction in MglA-mVenus and MglB-mCherry polar localization and partially reduced RomR-mCherry polar localization (Fig. 4E; Fig. S5C). We conclude that MglC^KRR^ is deficient in interacting with RomR and infer that the two positively charged KRR surface regions in the MglC dimer represent the interface to RomR.

### The α-helical RomR-C has three functions and represents the interface to MglC

RomR homologs comprise an N-terminal receiver domain of response regulators, an intrinsically disordered region (IDR), and an α-helical, negatively charged Glu-rich region at the C-terminus (RomR-C) (Fig. 5A)^30^. In BACTH assays, RomR-C interacts with MglC^36^. To examine whether RomR-C is the only interface to MglC, we generated RomR^1-368^ variants that lack RomR-C.

**Fig. 5.**
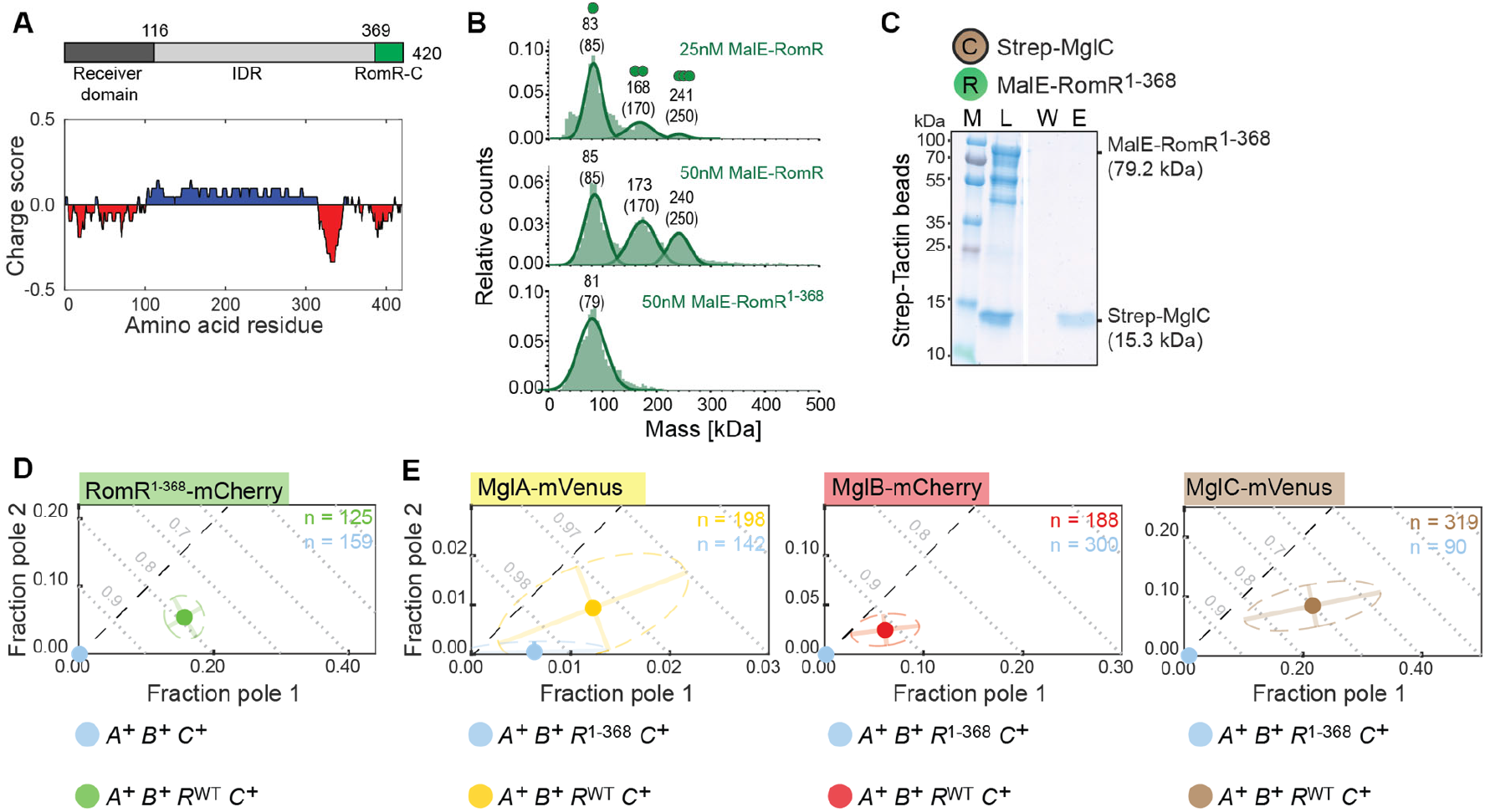
RomR-C has three functions and represents the interface for interaction with MglC1. A. Domain architecture and charge score of RomR. Numbering indicates amino acid positions. Charge score was calculated using a sliding window of 20 residues. B. MP analysis of MalE-RomR and MalE-RomR^1-368^. Molecular masses corresponding to the respective Gaussian fits are shown in kDa above the fittet curves. Calculated molecular masses of monomeric, dimeric and trimeric MalE-RomR and monomeric MalE-RomR^1-368^ are indicated in brackets together with symbols of oligomeric states. C. The RomR^1-368^ variant does not interact with MglC. Pull-down experiment was performed with Strep-MglC as bait on the indicated resin and presented as in Fig. 2. D. The RomR^1-368^-mCherry variant has strongly reduced polar localization. For comparison, RomR^WT^-mCherry is included (green dot). RomR^1-368^-mCherry was synthesized from the native locus. E. RomR^1-368^ causes strongly reduces polar localization of MglA, MglB and MglC. For comparison, the localization of the three fusion proteins is included in the presence of RomR^WT^ (yellow, red and brown dots). RomR^1-368^ was synthesized from the native locus. In D-E, data are presented as in Fig 1C.

First, using mass photometry (MP), we investigated the oligomeric structure of RomR. We detected MalE-RomR with masses matching well with monomers, dimers and trimers, while MalE-RomR^1-368^ was only detected at a mass matching monomers (Fig. 5B). Trimeric MalE-RomR was more prevalent at 50nM compared to 25nM (Fig. 5B) supporting that RomR forms up to trimers and begins to dissociate to dimers below 50nM. We conclude that RomR oligomerization depends on RomR-C and that the receiver domains and the IDRs do not interact. Moreover, based on quantitative immunoblot analysis, an *M. xanthus* cell contains ∼6000±2000 RomR molecules (Fig. S5E), resulting in a cellular RomR concentration of ∼2.5±0.8µM. We, therefore, suggest that RomR is predominantly present as a trimer *in vivo*.

In pull-down experiments, MalE-RomR^1-368^ did not detectably interact with Strep-MglC (Fig. 5C). Surprisingly, RomR^1-368^-mCherry polar localization in otherwise WT cells was strongly reduced (Fig. 5D; Fig. S5D). In the inverse experiments, RomR^1-368^, similar to the Δ*romR* mutation, caused strong reductions in the polar localization of MglA-mVenus, MglB-mCherry and MglC-mVenus (Fig. 5E; ^17, 30, 31^; Fig. S5D). We conclude that the negatively charged RomR-C has three functions: It is essential for RomR oligomerization, represents the RomR interface to MglC, and is critical for the polar localization of RomR.

### A structural model of the RomR/MglC/MglB complex

To gain structural insights into the RomR/MglC/MglB complex, we used structural information, our functional data and AlphaFold-Multimer structural predictions to model this complex. The AlphaFold-Multimer models of the MglB dimer and MglC dimer were predicted with high confidence and agreed well with the crystallographic structures^35, 40, 41^ (Fig. S7ABC), documenting the validity of the structural predictions.

A low-resolution structure of the MglC/MglB complex supports that one MglC dimer binds two MglB dimers^35^. In AlphaFold-Multimer models with the same stoichiometry, two MglB dimers are predicted with high accuracy to interact using their four-helix sides with the “lateral” edges of the two-helix side of the MglC dimer giving rise to an MglC_2_:(MglB_2_)_2_ complex (Fig. 6A; Fig. S7D). In this complex, Pymol-based analyses support that the R115 residues in the two KRK regions of an MglB dimer are involved in establishing contact with D26 and I28 of an MglC FDI region (Fig. 6A, inset). Thus, this structural model agrees with a 2:4 stoichiometry of the MglC/MglB complex and supports the experimental findings that the oppositely charged MglB KRK and MglC FDI surface regions interface.

**Fig. 6.**
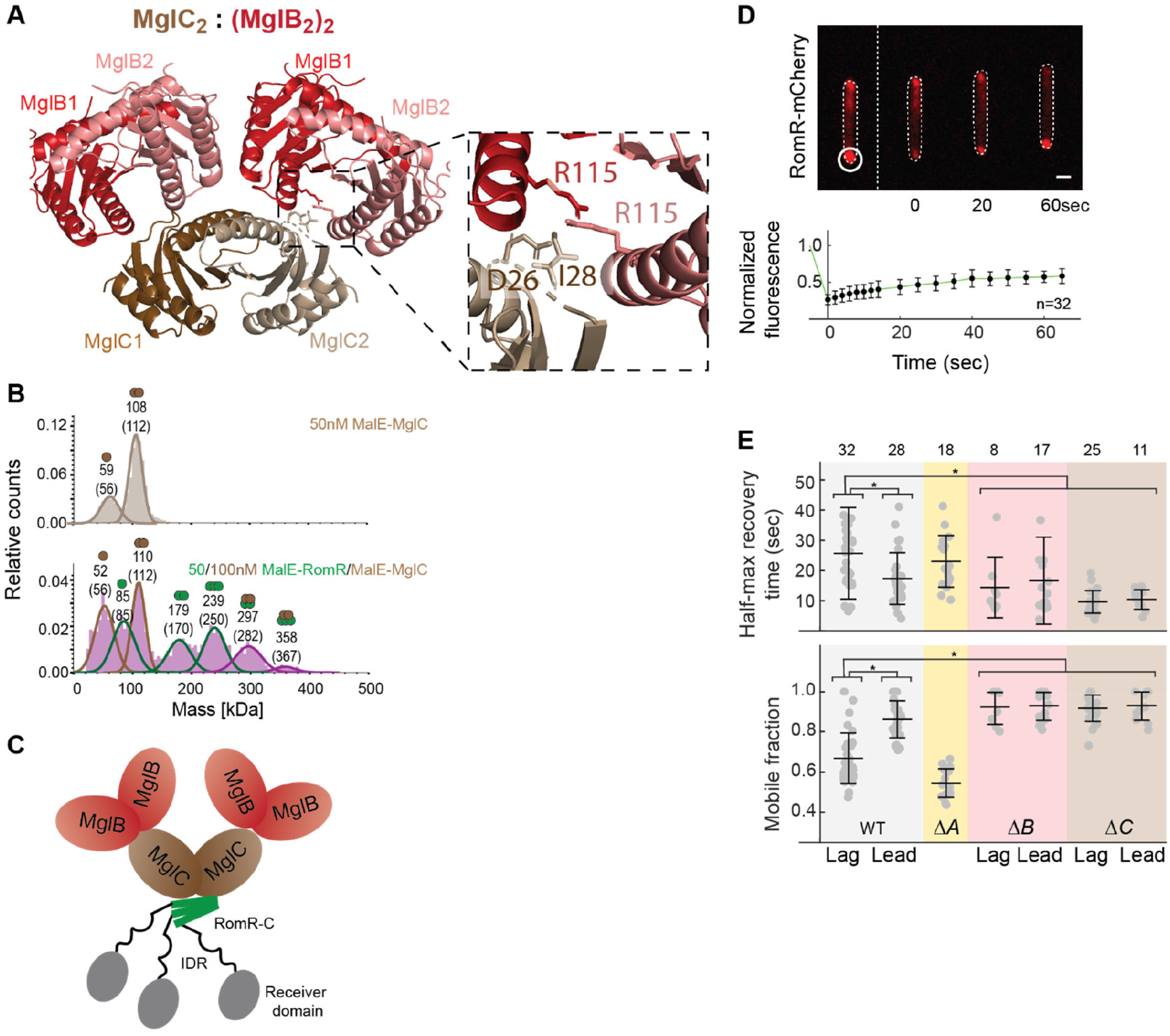
MglC and MglB stabilize polar RomR-mCherry binding. A. AlphaFold-Multimer structural model of MglC_2_:(MglB_2_)_2_ complex. Model rank 1 is shown. The inset shows R115 in each of the MglB protomers together with the D26 and I28 residues in a MglC protomer. B. MP analysis of MalE-MglC (top) and a mixture of MalE-RomR and MalE-MglC (bottom). Molecular masses corresponding to the respective Gaussian fits are shown in kDa above the fittet curves. Calculated molecular masses of monomeric and dimeric MalE-MglC, monomeric, dimeric and trimeric MalE-RomR, and MalE-RomR:MalE-MglC complexes with stoichiometries of 2:2 and 3:2 are indicated in brackets together with symbols of the oligomeric states. C. Schematic of the RomR/MglC/MglB complex with a 3:2:4 stoichiometry. D. Measurement of *in vivo* recovery kinetics of polar RomR-mCherry clusters in FRAP experiments. Upper panels, white circle indicates the bleached region of interest (ROI) at a lagging pole and the stippled line the bleaching event. Lower panel, normalized fluorescence intensity of the ROI before bleaching was set to 1.0. Colored dots indicate the mean and error bars STDEV. Dark lines show the recovery fitted to a single exponential. n, number of bleaching events at a lagging pole. Scale bar, 2μm. E. Summary of T_1/2_ and F_mob_. Cells were treated as in D with bleaching of clusters at the lagging (Lag) or leading (Lead) pole. Number of bleaching events listed above. Error bars, mean±STDEV. * *P*<0.05, two-sided Student’s t-test.

To determine the stoichiometry of the RomR/MglC complex, we used MP. We detected a MalE-MglC fusion protein with masses matching a monomer and dimer (Fig. 6B). In the presence of both MalE-MglC and MalE-RomR, we detected, in addition to the masses of the individual proteins, complexes with masses consistent with a RomR:MglC stoichiometry of 2:2 and 3:2 (Fig. 6B; see also Fig. 5B). To obtain structural insights into the RomR/MglC complexes, we attempted to generate AlphaFold-Multimer structural models of dimeric and trimeric RomR as well as of RomR_2_:MglC_2_ and RomR_3_:MglC_2_ complexes. However, none of these four complexes was predicted with high confidence. Altogether, our experimental data support that the MglC dimer can interact with dimeric and trimeric RomR and that the interface between MglC and RomR are represented by the oppositely charged KRR regions in the MglC dimer and RomR-C in the RomR dimer and trimer.

In total, these data support that a single MglC dimer is sandwiched between two MglB dimers and a RomR dimer or trimer, giving rise to a RomR:MglC:MglB complex with a 2:2:4 or a 3:2:4 stoichiometry. Because quantitative immunoblot analysis support that RomR is predominantly present as a trimer *in vivo*, we suggest that the dominant form of the RomR:MglC:MglB complex *in vivo* has a 3:2:4 stoichiometry (Fig. 6C).

### MglC and MglB decrease RomR-mCherry polar turnover

The structural model of the RomR/MglC/MglB complex sheds light on how RomR, MglC and MglB interact and how polar RomR recruits MglC, which recruits MglB. However, from this model, it is not clear how the positive RomR/MglC/MglB feedback is closed. We speculated that this loop could be closed if RomR would bind more stably to the poles in the RomR/MglC/MglB complex compared to RomR alone. To obtain a metric for the stability of RomR in polar clusters, we used Fluorescence Recovery after Photobleaching (FRAP) experiments in which polar RomR-mCherry clusters were bleached and half-maximal recovery time (T_1/2_) and the mobile fraction (F_mob_) used to assess RomR-mCherry turnover. In WT, RomR-mCherry at the lagging/leading pole dynamically exchanged with the cytoplasm with T_1/2_ of 25.7±15.2/17.3±8.6s, similar to previously published results^34^, and F_mob_ of 0.7±0.1/0.9±0.1 (Fig. 6DE). Thus, RomR-mCherry turnover is significantly lower at the lagging than at the leading pole. These observations agree with MglA-GTP at the leading pole engaging in a negative feedback to inhibit the RomR/MglC/MglB positive feedback. Consistently, in the non-motile Δ*mglA* mutant in which leading and lagging poles cannot be distinguished, T_1/2_ was increased, and F_mob_ decreased compared to the leading pole in WT (Fig. 6E). Importantly, in Δ*mglB* and Δ*mglC* cells, T_1/2_ and F_mob_ of RomR-mCherry at the two poles were similar, and the T_1/2_ values significantly lower and the F_mob_ values significantly higher than at the lagging pole in WT (Fig. 6E).

These observations support that MglB and MglC jointly reduce polar RomR-mCherry turnover, thus supporting that more stable polar binding of RomR in the presence of both MglC and MglB closes the RomR/MglC/MglB positive feedback.

### MglA-GTP breaks the MglC/MglB interaction

To dissect how MglA-GTP inhibits the positive RomR/MglC/MglB feedback, we hypothesized that MglA-GTP breaks the interaction between RomR/MglC, MglC/MglB or both. To this end, we performed pull-down experiments with Strep-MglC as bait. Strep-MglC pulled-down His_6_-MglB and MalE-RomR but not MglA-His_6_ in the presence of either the non-hydrolyzable GTP analogue GppNHp or GDP (Fig. 7AB; Fig S8). Intriguingly, in the presence of MglA-His_6_ loaded with GppNHp, Strep-MglC no longer pulled-down His_6_-MglB but still pulled-down His_6_-MglB in the presence of MglA-His_6_ loaded with GDP (Fig. 7A). By contrast, Strep-MglC pulled-down MalE-RomR in the presence of MglA-His_6_ loaded with either GppNHp or GDP (Fig. 7B). We conclude that MglA-GTP specifically inhibits the MglC/MglB interaction but not the MglC/RomR interaction.

**Fig. 7.**
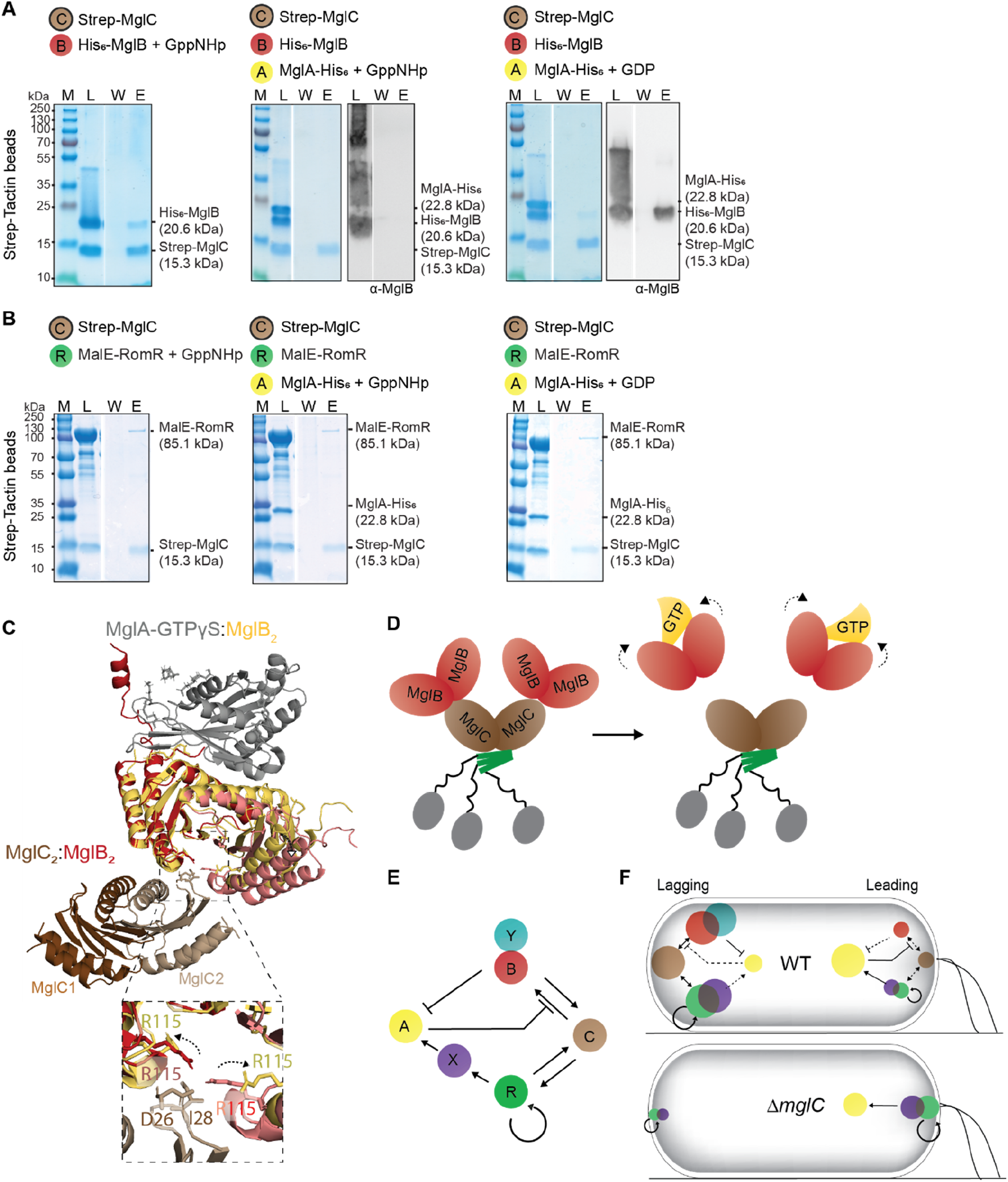
MglA-GTP breaks the interaction between MglC and MglB. A, B. MglA-GTP breaks the interaction between MglC and MglB. Pull-down experiments were performed with Strep-MglC as bait on the indicated resin as described in Fig. 2. MglA-His_6_ was preincubated with GppNHp or GDP (final concentration 40µM). All buffers contained 40µM GppNHp or GDP. In A, the SDS-PAGE gels were probed by immunoblotting with α-MglB antibodies. C. Crystallographic structure of MglA-GTPγS:MglB_2_ (pdb ID: 6izw^41^) superimposed on the AlphaFold-Multimer model of a MglC_2_:MglB_2_ complex (red/brown). Inset, R115 in each of the MglB protomers in the MglA-GTPγS:MglB_2_ complex (yellow) and the MglC_2_:MglB_2_ model (red) together with D26 and I28 (brown) in one of the MglC protomers. The arrows indicate the repositioning of R115 in the two complexes. For simplicity, the MglC dimer is shown to interact with only one MglB dimer. D. Schematic of the breaking of the MglB/MglC interaction by MglA-GTP. Bent arrows indicate the conformational change in the MglB dimers upon binding of MglA-GTP. E. Regulatory interactions that establish and maintain front-rear polarity in *M. xanthus*. F. Different interactions between the polarity proteins dominate at the leading and lagging poles in WT and the Δ*mglC* mutant. Full arrows show locally strong interactions, dashed arrows show interactions that are locally suppressed. Color code as in E.

To understand the basis for MglA-GTP inhibition of the MglC/MglB interaction, we compared the MglA-GTPɣS:MglB_2_ crystallographic structure to the MglC_2_:(MglB_2_)_2_ AlphaFold-Multimer model. We identified significant conformational differences in the MglB dimers in the two complexes. Specifically, the MglB_2_ four-helix side is in a more open state when complexed with MglA-GTPɣS than with MglC_2_ (Fig. 7C). Consequently, the two R115 residues in the MglB KRK regions are likely positioned in such a way in the complex with MglA-GTPɣS that they cannot interact with D26 and I28 in the MglC FDI region (Fig. 7CD; see also Fig. 6A).

## Discussion

Here, we identify MglC as a critical component of the polarity module for switchable front-rear polarity in *M. xanthus*. We demonstrate that the previously proposed RomR/MglB positive feedback incorporates and depends on MglC. These three proteins form a heteromeric RomR/MglC/MglB complex in which MglC is sandwiched between RomR and MglB. *In vivo*, they establish the RomR/MglC/MglB positive feedback that results in the colocalization of the RomR/RomX GEF and MglB/RomY GAP at high concentration at the lagging pole (Fig. 7E and F, upper panel). Moreover, we demonstrate that the previously reported inhibitory effect of MglA-GTP on the RomR/MglB positive feedback is the result of MglA-GTP breaking the MglC/MglB interaction without interfering with the RomR/MglC interaction in the RomR/MglC/MglB positive feedback (Fig. 7E and F, upper panel). By way of this inhibitory effect, MglA-GTP at the leading pole limits the accumulation of the other polarity regulators at this pole. By engaging in these interactions, MglC stimulates polar localization of the remaining polarity proteins and is also key to enabling dynamic inversion of polarity in response to Frz signaling.

*In vitro* observations together with an AlphaFold-Multimer structural model of the MglC/MglB complex and *in vivo* experiments, support that the RomR:MglC:MglB complex has a 3:2:4 stoichiometry. Specifically, our data support that the negatively charged α-helical RomR-C interacts with the two juxtaposed positively charged MglC KRR surface regions in the dimer, and that each of the two negatively charged FDI surface regions in the MglC dimer interface with the positively charged KRR surface regions in a MglB dimer. These interactions between oppositely charged surface regions allow polar RomR to recruit MglC, which recruits MglB. FRAP experiments *in vivo* demonstrated that MglC and MglB enable more stable polar RomR occupancy. Based on these findings, we infer that the RomR/MglC/MglB positive feedback for polar localization involves direct recruitment *via* the RomR→MglC→MglB interactions. These interactions stabilize polar RomR binding, thereby closing the positive feedback. Because neither RomR, MglC, nor RomR/MglC has measurable GAP activity or measurably affects GAP activity by MglB/RomY and MglB, we infer that one role of MglC is to connect MglB and RomR to establish the positive feedback.

*In vitro*, MglA-GTP breaks the MglC/MglB interaction in the RomR/MglC/MglB complex without interfering with the RomR/MglC interaction. A comparison of the solved structure of the MglA-GTPγS:MglB_2_ complex with an AlphaFold-Multimer model of the MglC_2_:(MglB_2_)_2_ complex supports that MglA-GTP breaks the MglC/MglB interaction using an allosteric mechanism. Specifically, MglA-GTP by binding to the two-helix side of a MglB homodimer induces a conformational change that alters the four-helix side the MglB homodimer, thereby breaking the interaction between the MglC FDI and MglB KRK interfaces. Thus, the second role of MglC is to enable the inhibitory effect of MglA-GTP on the RomR/MglC/MglB positive feedback *in vivo*.

Our study raises several intriguing questions for future research avenues regarding the proteins of the polarity module. First, RomR-C has three functions: It interacts not only with MglC but also mediates oligomerization with dimer and trimer formation and is also essential for RomR polar localization. *In vivo* quantification of the RomR concentration suggests that trimeric RomR is the active form *in vivo*; however, it is unknown whether dimeric RomR has a function. Similarly, it is not known how Rom-C brings about the polar localization of RomR, and how RomR stimulates its polar binding. Second, experimental evidence and AlphaFold-Multimer structural models support that MglB binds its co-GAP RomY with low affinity on the two-helix side^28^. We, therefore, suggest that the RomR/MglC/MglB complex at the lagging pole also contains RomY forming a RomR/MglC/MglB/RomY complex. Third, RomR interacts with RomX to generate the polarly localized Rom/RomX GEF complex. While this complex’s structural details are unknown, they raise the possibility that the RomR/MglC/MglB complex may also include RomX. The complexes formed will be addressed in future work.

The Δ*mglC* mutant resembles WT concerning unidirectional motility but is less sensitive to Frz signaling, supporting that the ultimate function of MglC is to establish sensitivity to Frz signaling, thereby enabling polarity inversions. The two output response regulators of the Frz system, FrzX and FrzZ, act on the polarity module by unknown mechanisms to enable polarity inversions^30, 31, 33, 34^. The observation that the Δ*mglC* mutant still responds to high levels of Frz signaling argues that MglC is not the downstream molecular target of the Frz system but enables Frz responsiveness by a different mechanism. As predicted by the model for polarity establishment (Fig. 7EF, upper panels), neither MglC nor the RomR/MglC/MglB positive feedback is important for MglA localization at the leading pole.

Instead, in the absence of MglC, and therefore, the RomR/MglC/MglB positive feedback, the highest polar concentration of the RomR/RomX complex colocalizes with MglA at the leading pole (Fig. 7F, lower panel). In the Δ*mglC* mutant, RomR/RomX and MglA polar localization is driven by RomR stimulating its own polar binding in a positive feedback and then recruiting RomX and MglA (Fig. 7F, lower panel). Thus, in this configuration, the polarity module is less sensitive to the Frz system, while front-rear polarity is robustly maintained. Based on theoretical arguments, we previously argued that the configuration with a high concentration of the RomR/RomX GEF at the lagging pole would allow for the rapid accumulation of MglA-GTP at this pole in response to Frz signaling. We, therefore, suggest that the spatial configuration of the polarity proteins in the Δ*mglC* mutant makes it less sensitive to Frz signaling because there is too little RomR/RomX GEF at the lagging pole to recruit MglA-GTP during reversals. Thus, the RomR/MglC/MglB positive feedback resulting in the peculiar colocalization of the RomR/RomX GEF and MglB/RomY GAP at the lagging pole in WT serves two purposes: First, the GAP activity displaces MglA-GTP from this pole to enable unidirectional translocation; and, second, the GEF activity is necessary to provide the system with the ability to rapidly and efficiently invert polarity. In other words, an important role of MglC and the RomR/MglC/MglB positive feedback is to establish the configuration of the polarity proteins that confer the polarity module with responsiveness to the Frz system.

Returning to the question raised in the introduction, i.e. why different network designs have been selected for in various polarity-regulating networks with functionally equivalent outcomes, in the *S. cerevisiae* polarity system that establishes the single Cdc42 cluster, the positive feedback is centred on Cdc42 and the Cdc24 GEF^9^. Therefore, once the Cdc42 cluster is established, this polarity is stably maintained, and the decay of a nascent bud site or the formation of competing bud sites is efficiently avoided. In the Δ*mglC* mutant, RomR, by stimulating its own polar binding in a positive feedback, brings about RomR/RomX and MglA polar localization at the same pole (Fig. 7F, lower panel). This design is conceptually similar to the yeast system driving Cdc42 cluster formation. Thus, while the network designs of the *M. xanthus* and the *S. cerevisiae* polarity systems enable the formation of a single MglA/Cdc42 cluster, the different wirings can be rationalized as the *M. xanthus* polarity module being part of a spatial toggle switch that is optimal for stable polarity as well for rapid polarity inversions. By contrast, the *S. cerevisiae* system is optimized to provide stable polarity.

In principle, it would seem that the RomR/MglC/MglB positive feedback could have been established by RomR interacting directly with MglB, raising the question of the advantage of incorporating MglC into the RomR/MglC/MglB positive feedback. The roadblock domain protein family is ancient, abundantly present in all domains of life, and often involved in regulating GTPase activity^37, 43-45^. Interestingly, the Rag GTPases of the mTOR pathway are composed of a small GTPase domain and a C-terminal roadblock domain and form heterodimers using their roadblock domains^46^. These heterodimers are recruited to lysosomes by the Ragulator complex, which contains two roadblock heterodimers that interact head-to-tail forming a tetrameric complex^46^. The Rag GTPase/Ragulator interaction occurs via the roadblock domains, resulting in three layers of heterodimeric roadblock domains^46^. Intriguingly, Rag heterodimers’ GTP/GDP state allosterically regulates their binding to Ragulator by tuning the interaction between pairs of roadblock heterodimers^47, 48^. This mechanism is conceptually remarkably similar to the GTP/GDP state of MglA regulating the interaction between the MglC/MglB homodimers, supporting that this regulatory mechanism is evolutionary conserved. We suggest that the presence of MglC in the RomR/MglC/MglB positive feedback reflects an ancient regulatory mechanism in which the GTP/GDP state of a partner GTPase can modulate the interaction between pairs of roadblock dimers.

## Supporting information

All supplementary information

## Acknowledgements

We thank Dr. Anna McLoon for the generation of the α-MglC antibodies. The Max Planck Society generously supported this work.

## Data Availability

The data that support the findings of this study are included in the manuscript or in the Supplementary Information.

## Author contributions

Conceptualization: Luís António Menezes Carreira, Lotte Søgaard-Andersen

Experimental work: Luís António Menezes Carreira, Dobromir Szadkowski, Stefano Lometto

Analysis of experimental data: Luís António Menezes Carreira, Stefano Lometto

Writing – original draft: Luís António Menezes Carreira, Lotte Søgaard-Andersen

Writing – editing of draft: Luís António Menezes Carreira, Dobromir Szadkowski, Stefano

Lometto, Georg K.A. Hochberg, Lotte Søgaard-Andersen

Supervision: Georg K.A. Hochberg, Lotte Søgaard-Andersen,

Funding acquisition: Georg K.A. Hochberg, Lotte Søgaard-Andersen

## Methods

### Cell growth and construction of strains

Strains, plasmids and primers used in this work are listed in Supplementary Table 1, 2 and 3, respectively. All *M. xanthus* strains are derivatives of the DK1622 WT strain^49^. *M. xanthus* was grown at 32°C in 1% CTT broth^50^ or on 1.5% agar supplemented with 1% CTT and kanamycin (50µg mL^-1^) or oxytetracycline (10µg mL^-1^) as appropriate. In-frame deletions were generated as described^51^. Plasmids were introduced in *M. xanthus* by electroporation and integrated by homologous recombination at the endogenous locus or at the *mxan18-19* locus or by site-specific recombination at the Mx8 *attB* site. All in-frame deletions and plasmid integrations were verified by PCR. Plasmids were propagated in *Escherichia coli* TOP10 (F^-^, *mcrA*, ∆(*mrr-hsd*RMS-*mcr*BC), φ80*lac*Z∆M15, ∆*lac*X74, *deo*R, *rec*A1, *ara*D139, ∆(*ara-leu*)7679, *gal*U, *gal*K, *rps*L, *end*A1, *nup*G). *E. coli* was grown in LB or on plates containing LB supplemented with 1.5% agar at 37°C with added antibiotics when appropriate^52^. All DNA fragments generated by PCR were verified by sequencing.

### Motility assays and determination of reversal frequency

Population-based motility assays were done as described^53^. Briefly, *M. xanthus* cells from exponentially growing cultures were harvested at 4000× *g* for 10min at room temperature (RT) and resuspended in 1% CTT to a calculated density of 7×10^9^ cells mL^-1^. 5µL aliquots of cell suspensions were placed on 0.5% agar plates supplemented with 0.5% CTT for T4P-dependent motility and 1.5% agar plates supplemented with 0.5% CTT for gliding motility and incubated at 32°C. After 24h, colony edges were visualized using a Leica M205FA stereomicroscope and imaged using a Hamamatsu ORCA-flash V2 Digital CMOS camera (Hamamatsu Photonics). For higher magnifications of cells at colony edges on 1.5% agar, cells were visualized using a Leica DMi8 inverted microscope and imaged with a Leica DFC9000 GT camera.

Individual cells were tracked as described^29^. Briefly, for T4P-dependent motility, 5µL of exponentially growing cultures were spotted into a 24-well polystyrene plate (Falcon). After 10min at RT, cells were covered with 500µL 1% methylcellulose in MMC buffer (10mM MOPS (3-(*N*-morpholino) propanesulfonic acid) pH 7.6, 4mM MgSO_4_, 2mM CaCl_2_), and incubated at RT for 30min. Subsequently, cells were visualized for 10min at 20sec intervals at RT using a Leica DMi8 inverted microscope and a Leica DFC9000 GT camera. Individual cells were tracked using Metamorph 7.5 (Molecular Devices) and ImageJ 1.52b^54^ and then the speed of individual cells per 20sec interval as well as the number of reversals per cell per 10min calculated. For gliding, 5µL of exponentially growing cultures were placed on 1.5% agar plates supplemented with 0.5% CTT, covered by a cover slide and incubated at 32°C. After 4 to 6h, cells were observed for 15min at 30sec intervals at RT as described, speed per 30sec interval as well as the number of reversals per 15min calculated.

### Immunoblot analysis

Immunoblot analysis was done as described^52^. Rabbit polyclonal α-MglA^27^, α-MglB^27^, α-RomR^32^, α-PilC^55^, PilO^56^ and α-MglC antibodies were used together with horseradish peroxidase-conjugated goat anti-rabbit immunoglobulin G (Sigma) as a secondary antibody. Mouse anti-GFP antibodies (Sigma) were used together with horseradish peroxidase conjugated sheep anti-mouse immunoglobulin G (GE Healthcare) as a secondary antibody. To generate rabbit polyclonal α-MglC antibodies, His_6_-MglC was purified as described (see below) and used for immunization as described^52^. Blots were developed using Luminata Crescendo Western HRP substrate (Millipore) and visualized using a LAS-4000 luminescent image analyzer (Fujifilm). Proteins were separated by SDS-PAGE as described ^52^.

### Fluorescence microscopy

For fluorescence microscopy, exponentially growing cells were placed on slides containing a thin pad of 1% SeaKem LE agarose (Cambrex) with TPM buffer (10mM Tris-HCl pH 7.6, 1mM KH_2_PO_4_ pH 7.6, 8mM MgSO_4_) and 0.2% CTT, and covered with a coverslip. After 30min at 32°C, cells were visualized using a temperature-controlled Leica DMi8 inverted microscope and phase contrast and fluorescence images acquired using a Hamamatsu ORCA-flash V2 Digital CMOS camera. For time-lapse recordings, cells were imaged for 15min using the same conditions. To induce expression of genes from the vanillate inducible promoter^57^, cells were treated as described in the presence of 300μM vanillate and imaged for 6h. To precisely quantify the localization of fluorescently-labelled proteins, we used an established analysis pipeline^17^ in which the output for each cell is total cellular fluorescence, the fractions of fluorescence in clusters at each pole, and the mean fraction of total polar fluorescence. For calculating mean fraction of total polar fluorescence cells with and without clusters were included. The quantification of fluorescence signals is included in Supplementary Table 4.

### Image analysis

Microscope images were processed with Fiji^58^ and cell masks determined using Oufti^59^ and manually corrected when necessary. Fluorescence was quantified in Matlab R2020a (The MathWorks) using custom scripts as described ^17^.

### *In vivo* fluorescence recovery after photobleaching (FRAP)

FRAP experiments were performed as described^60^ with a temperature-controlled Nikon Ti-E microscope with Perfect Focus System and a CFI PL APO 100x/1.45 Lambda oil objective at 32°C with a Hamamatsu Orca Flash 4.0 camera using NIS Elements AR 2.30 software (Nikon) in the dark. Photobleaching was performed using a single circular shaped region with 20% laser power (561nm) and a 500µsec dwelling time. For every image, integrated fluorescence intensities of a whole cell and the bleached region of interest (ROI), were measured. After background correction, the corrected fluorescence intensity of the bleached ROI was divided by total corrected cellular fluorescence, correcting for bleaching effects during picture acquisition. Cell segmentation and background correction was performed with Oufti. This normalized fluorescence was correlated to the initial fluorescence in the ROI. The mean relative fluorescence of several cells was plotted as a function of time. The recovery rate for a given fluorescent protein was determined by fitting the plotted data to a single exponential equation with Matlab R2020a (The MathWorks).

### Protein purification

All proteins were expressed in *E. coli* Rosetta 2(DE3) (F^−^ *ompT hsdS*_B_ (r_B_ ^−^ m_B_^−^) *gal dcm* (DE3 pRARE2) at 18°C or 37°C. To purify His_6_-tagged proteins, Ni-NTA affinity purification was used. Briefly, cells were washed in buffer A (50mM Tris-HCl pH 7.5, 150mM NaCl, 10mM imidazole, 5% glycerol, 5mM MgCl_2_) and resuspended in lysis buffer A (50 mL of wash buffer A supplemented with 1mM DTT, 100µg ml^−1^ phenylmethylsulfonylfluoride (PMSF), 10U ml^−1^ DNase 1 and Complete Protease Inhibitor Cocktail Tablet (Roche)). Cells were lysed by sonication and cell debris was removed by centrifugation (48,000× *g*, 4°C, 30min) and filtration through a 0.45µm filter (Sarsted). The cleared cell lysate was loaded onto a 5mL HiTrap Chelating HP column (Cytiva) preloaded with NiSO_4_ as described by the manufacturer and pre-equilibrated in buffer A. The column was washed with 20 column volumes of column wash buffer (buffer A with 20mM imidazole). Proteins were eluted with elution buffer (buffer A with 500mM imidazole) using a linear imidazole gradient from 20 to 500mM. Fractions containing purified His_6_-tagged proteins were combined and loaded onto a HiLoad 16/600 Superdex 75 pg (GE Healthcare) gel filtration column that was equilibrated with buffer 1 (50mM Tris-HCl pH 7.5, 150mM NaCl, 1mM DTT, 5mM MgCl_2_, 5% glycerol). Fractions containing His_6_-tagged proteins were pooled, frozen in liquid nitrogen and stored at −80°C.

To purify MalE-tagged proteins (MalE-RomR and MalE-MglC), maltose-binding protein (MBP) affinity purification was used. Briefly, cells were washed in buffer B (50mM Tris-HCl pH 7.5, 150mM NaCl, 1mM EDTA, 1mM DTT) and resuspended in 50mL lysis buffer B (50mL buffer B supplemented with PMSF 100µg mL^−1^, DNase 1 10U mL^−1^ and Complete Protease Inhibitor Cocktail Tablet (Roche)). Cells were lysed and cleared cell lysates prepared as described and loaded onto a 5mL MBPTrapHP (Cytiva) column equilibrated with buffer B. The column was washed with 20 column volumes of buffer B. Proteins were eluted with elution buffer B (buffer B with 10mM maltose). Eluted fractions containing MalE-RomR or MalE-MglC were loaded onto a 5 mL HiTrap Q HP ion exchange column (Cytiva) equilibrated with buffer C (50mM Tris-HCl pH 7.5, 50mM NaCl, 5mM MgCl_2_, 1mM DTT, 5% glycerol). The column was washed with 20 column volumes of buffer C. MalE-RomR or MalE-MglC were eluted with buffer C using a linear gradient of NaCl from 50 to 500mM. Fractions containing MalE-RomR or MalE-MglC were loaded onto a HiLoad 16/600 Superdex 200 pg (GE Healthcare) gel filtration column that was equilibrated with buffer 1. Fractions with MalE-RomR or MalE-MglC were pooled, frozen in liquid nitrogen and stored at −80°C.

To purify Strep-tagged proteins, biotin affinity purification was used. Briefly, cells were washed in buffer C (100mM Tris-HCl pH 8.0, 150mM NaCl, 1mM EDTA, 1mM DTT) and resuspended in lysis buffer C (50mL of wash buffer C supplemented with 100µg mL^−1^ PMSF, 10U mL^−1^ DNase 1 and Complete Protease Inhibitor Cocktail Tablet (Roche)). Cells were lysed and cleared lysate prepared as described and loaded onto a 5mL Strep Trap HP (Cytiva) column, equilibrated with buffer C. The column was washed with 20 column volumes of buffer C. Protein was eluted with elution buffer C (buffer C with 2.5mM desthiobiotin). Elution fractions containing Strep-tagged proteins were loaded onto a HiLoad 16/600 Superdex 75 pg (GE Healthcare) gel filtration column that was equilibrated with buffer 1. Fractions with Strep-tagged proteins were pooled, frozen in liquid nitrogen and stored at −80°C.

### Pull-down experiments

To test for interactions with MglA, MglB and RomR, Strep-MglC (final concentration 10µM) was incubated with MglA-His_6_, His_6_-MglB or MalE-RomR (final concentration 10µM) in buffer 1 (50mM Tris-HCl pH 7.5, 150mM NaCl, 1mM DTT, 5mM MgCl_2_, 5% glycerol) for 30min at RT. Subsequently, 10µL of Strep-Tactin MagStrep’ type3’ XT beads (IBA Lifesciences) previously equilibrated with buffer 1 was added for 30min at RT. Then beads were washed 10 times with 1mL buffer 1. Proteins were eluted with 200µL elution buffer (100mM Tris-HCl pH 8.0, 150mM NaCl, 1mM EDTA, 50mM biotin). To test for interactions with MglC, MglA and MglB, MalE-RomR (final concentration 10µM) was incubated with Strep-MglC, MglA-His_6_ and/or His_6_-MglB (final concentration 10µM) in buffer 1 for 30min at RT. Subsequently, the mixture was added to 200µL of Amylose Resin, previously equilibrated with buffer 1, and incubated for 30min at RT. The resin was then washed 10 times with 1mL buffer 1. Proteins were eluted with 200µL elution buffer (100mM Tris-HCl pH 8.0, 150mM NaCl, 1mM EDTA, 10mM amylose). To test for interactions with MglC, MglA and RomR, His_6_-MglB (final concentration 10µM) was incubated with Strep-MglC, MglA-His_6_ and/or MalE-RomR (final concentration 10 µM) in buffer 1 for 30min at RT. Subsequently, 20µL of Amintra Nickel Magnetic beads (Expedeon), previously equilibrated with buffer 1, was added to the mixture and incubated for 30min at RT. Beads were then washed 10 times with 1mL buffer 2 (buffer 1 with 50mM imidazole). Proteins were eluted with 200µL elution buffer (buffer 1 with 500mM imidazole). In experiments involving MglA-His_6_, MglA-His_6_ (final concentration 10µM) was preloaded with GTP, GDP or GppNHp (final concentration 40µM) for 30min at RT in buffer 1 and all buffers contained 40µM of the relevant nucleotide.

### GTPase assays

GTP-hydrolysis by MglA-His_6_ was measured using a continuous, regenerative coupled GTPase assay^61^ in reaction buffer (50mM Tris-HCl pH 7.5, 150mM NaCl, 5% glycerol, 1mM DTT, 7.5mM MgCl_2_) supplemented with 495µM NADH (Sigma), 2mM phosphoenolpyruvate (Sigma), 18-30U mL^-1^ pyruvate kinase (Sigma) and 27-42 U mL^-1^ lactate dehydrogenase (Sigma). For all assays, MglA-His_6_ (final concentration 2µM) was preloaded with GTP (final concentration 3.3mM) for 30min at RT in reaction buffer. In parallel, MglB was preincubated with Strep-MglC, MalE-RomR and/or Strep-RomY for 10min at RT in reaction buffer. GTPase reactions were performed in 96-well plates (Greiner Bio-One) and initiated by adding His_6_-MglB, Strep-MglC, MalE-RomR and/or Strep-RomY to the MglA/GTP mixture. Final concentration, MglA-His_6_: 2µM, His_6_-MglB: 4µM, Strep-MglC: 4µM, MalE-RomR: 2µM, Strep-RomY: 2µM, GTP: 1mM. Absorption was measured at 340nm for 60min at 37°C using an Infinite M200 Pro plate-reader (Tecan) and the amount of hydrolyzed GTP per hour per molecule of MglA-His_6_ calculated. For each reaction, background subtracted GTPase activity was calculated as the mean of three technical replicates.

### Mass photometry (MP)

MP was performed using a TwoMP mass photometer (Refeyn Ltd, Oxford, UK). Data acquisition was performed using AcquireMP (Refeyn Ltd. v2.3). MP movies were recorded at 1 kHz, with exposure times varying between 0.6 and 0.9 ms, adjusted to maximize camera counts while avoiding saturation. Microscope slides (1.5 H, 24×50mm, Carl Roth) and CultureWellTM Reusable Gaskets were cleaned with three consecutive rinsing steps of double-distilled H_2_O and 100% isopropanol and dried under a stream of pressurized air. For measurements, gaskets were assembled on coverslips and placed on the stage of the mass photometer with immersion oil. Assembled coverslips were held in place using magnets. For measurements, gasket wells were filled with 10µL of 1× phosphate-buffered saline (137mM NaCl, 2.7mM KCl, 8mM Na_2_HPO_4_, 2mM KH_2_PO_4_) to enable focusing of the glass surface. After focusing, 10µL sample were added, rapidly mixed while keeping the focus position stable and measurements started. MP contrast values were calibrated to molecular masses using an in-house standard. For each sample, three separate measurements were performed. The data were analyzed using the DiscoverMP software (Refeyn Ltd, v. 2022 R1). MP image analysis was done as described^62^.

### AlphaFold model generation

AlphaFold-multimer structure prediction was done with the ColabFold pipeline^63-65^. ColabFold was executed with default settings where multiple sequence alignments were generated with MMseqs2^66^ and HHsearch^67^. The ColabFold pipeline generates five model ranks. Predicted Local Distance Difference Test (pLDDT) and alignment error (pAE) graphs were generated for each rank with a custom Matlab R2020a (The MathWorks) script. Ranking of the models was performed based on combined Plddt and pAE values, with the best-ranked models used for further analysis and presentation. Per residue model accuracy was estimated based on pLDDT values (>90, high accuracy; 70-90, generally good accuracy; 50-70, low accuracy; <50, should not be interpreted)^64^. Relative domain positions were validated by pAE^64^. Only models of the highest confidence, based on combined pLDDT and pAE values, were used for further investigation. For all models, sequences of full-length proteins were used.

### Bioinformatics

Sequence alignments were generated using ClustalOmega^68^ with default parameters and alignments were visualized with Jalview^69^. Protein domains were identified using Interpro^70^. Charge score was calculated using the Protein-sol tool^71^. Structural alignments and calculation of electrostatic surface potential were done in Pymol (The PyMOL Molecular Graphics System, Version 1.2r3pre, Schrödinger, LLC).

### Statistics

Statistics were performed using a two-tailed Student’s *t*-test for samples with unequal variances.

