## Supplementary material for "Molecular basis and design principles of a system for switchable front-rear polarity and directional migration": All supplementary information

This file includes

- Supplementary Figure 1-8
- Supplementary Tables 1-4
- Supplementary References

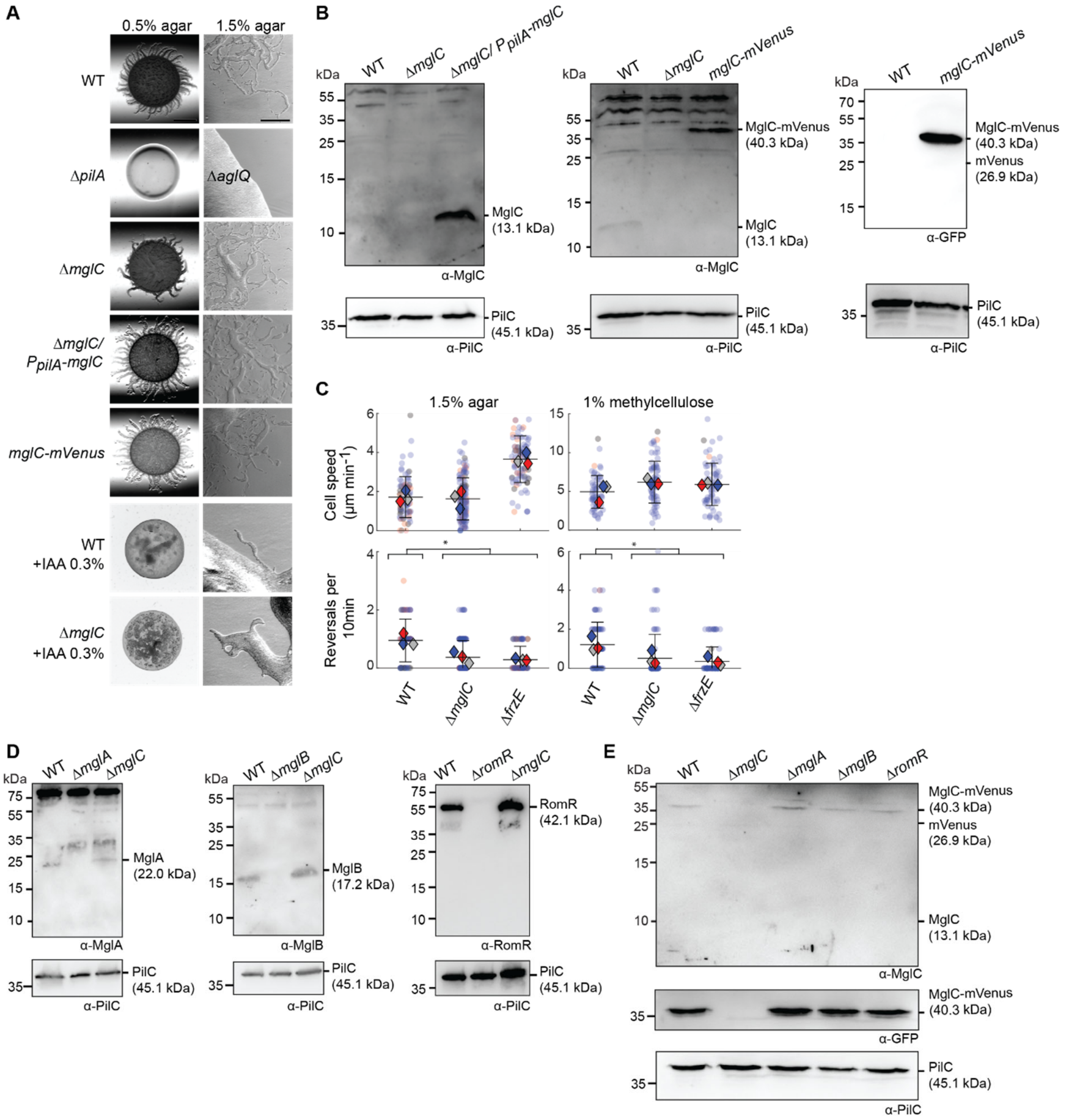


<u>Sup. Fig. 1. MglC is important for the correct reversal frequency</u>

A. MglC is important for T4P-dependent motility and gliding. Cells were incubated on 0.5% agar/0.5% CTT and 1.5% agar/0.5% CTT to score T4P-dependent and gliding motility, respectively and in the absence or presence of IAA. Scale bars, 1mm (0.5% agar), 500µm (1.5% agar, left). In the Δ*mglC*/P*pilA*-*mglC* complementation strain, *mglC* was expressed from the *pilA* promoter on a plasmid integrated in a single copy at the Mx8 *attB* site. *mglC-mVenus* is integrated at the native site. T4P-dependent motility and gliding motility is favored on 0.5% and 1.5% agar, respectively [1]. T4P-dependent motility is characterized by the formation of flares at the edge of colonies and gliding by the presence of single cells at the colony edge.

B. Immunoblot analysis of MglC and MglC-mVenus accumulation. Lysates from the same number of cells were separated by SDS-PAGE and probed with α-MglC or α-GFP antibodies and after stripping with α-PilC antibodies (loading control). Calculated molecular masses of the indicated proteins are indicated on the right and molecular weight masses of markers on the left.

C. MglC is important for the correct reversal frequency. Speed (upper panels) and reversals (lower panels) of single cells moving by gliding motility (1.5% agar, left) or T4P-dependent motility (covered with 1% methylcellulose, right) were determined. Data are shown for three biological replicates. Individual data points are shown in red, blue and grey and the corresponding mean (diamond) in the same color. Error bars indicate standard deviation (STDEV). >20 cells were analyzed per replicate. In all panels, * $P<0.01$, two-sided Student's *t*-test.
D, E. Immunoblot analysis of MglA, MglB, RomR and MglC-mVenus accumulation. Immunoblot analysis was done as in Fig. S1B.

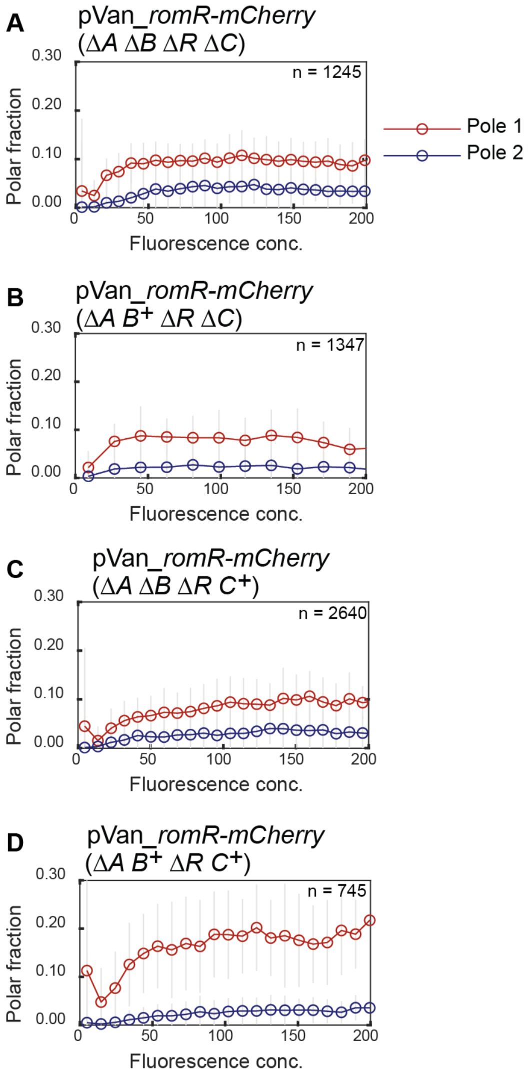


Sup. Fig. 2. MglC is not important for cooperative RomR polar accumulation

A-D. Induction of *romR-mCherry* expression from the vanillate inducible promoter in Δ*mglA* Δ*mglB* Δ*romR* Δ*mglC* cells (A), Δ*mglA*Δ*romR* Δ*mglC* cells (B), Δ*mglA* Δ*mglB* Δ*romR* cells (C) and Δ*mglA* Δ*romR* cells (D). All strains contain the Δ*aglQ* Δ*frzE* mutations to allow monitoring of the same cells for 6h and to reduce Frz-dependent polarity inversions. n, number of individual cells observed. In the diagrams, the poles with the highest and lowest polar fraction of fluorescence are defined as pole 1 and pole 2, respectively. Cells were grown in the absence of vanillate and then placed on an agar surface containing 300µM vanillate and imaged for 6h. Cells from all frames of the time-lapse recordings were pooled and binned according to their fluorescence concentration (total cellular fluorescence divided by cell area). Plotted are the mean ± 1 STDEV of all cells within each bin. Each bin contain data from at least five cells. Data in C and D are from[2]. The shape of the polar accumulation curves provides evidence for positive cooperativity in RomR-mCherry polar localization [2]: In the absence of any cooperativity, individual RomR-mCherry molecules would localize independently of one another, such that the fraction at each pole should be constant and independent of concentration. Instead, the fractions of RomR-mCherry at both poles increase with fluorescence concentration, supporting that RomR-mCherry self-recruits or stabilizes its polar accumulation.

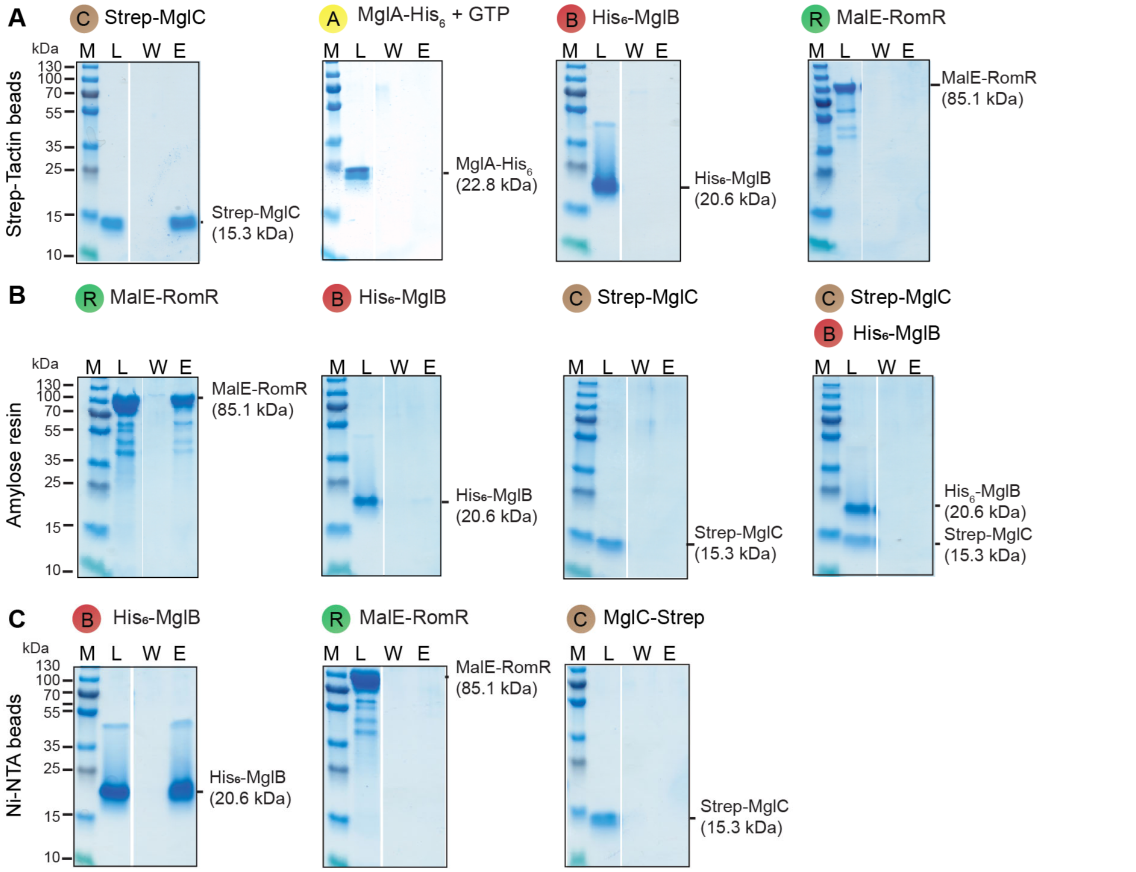


<u>Sup. Fig. 3. *In vitro* pull-down experiments with MglA-$His_6$, $His_6$-MglB, Strep-MglC and MalE-RomR</u>

A-C. Control experiments to the experiments shown in Fig. 2. Proteins were mixed as indicated at final concentrations of 10µM and applied to the indicated matrices. Matrices were washed and bound proteins eluted. In all panels, the bait protein is indicated by the black circle. In experiments with MglA-$His_6$, the protein was preloaded with GTP and all buffers contained 40µM GTP. Equivalent volumes of the load (L), last wash (W) and elution fraction (E) were separated on the same SDS-PAGE gel and stained with Coomassie Brilliant Blue. Gap between lanes indicates lanes deleted for presentation purposes. Calculated molecular masses of the indicated proteins are indicated on the right. Molecular weight markers (M) are indicated on the left.

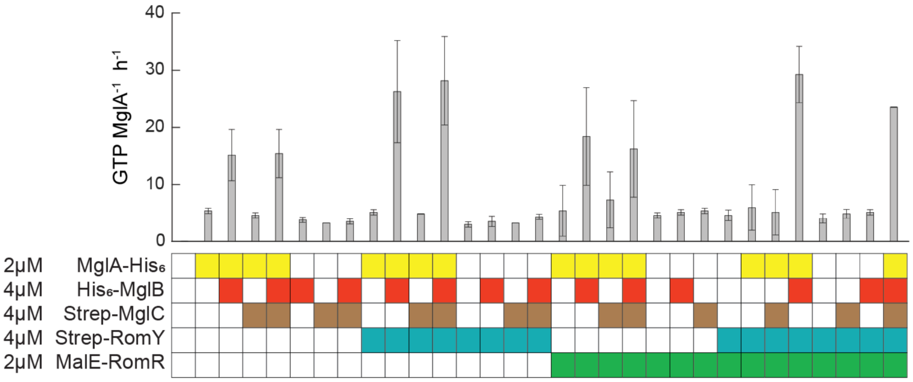


<u>Sup. Fig. 4. RomR, MglC and RomR/MglC do not have MglA GAP activity and do not affect MglB and MglB/RomY GAP activity.</u>
The indicated proteins at the indicated concentrations were mixed and GTPase activity measured in a continuous, regenerative coupled GTPase assay after 1h of incubation. Boxes below diagrams indicate the presence (colored box) or absence (white box) of a protein. MglA-$His_6$ was preloaded with GTP (final concentration 3.3mM). Error bars, mean±STDEV from three technical replicates.

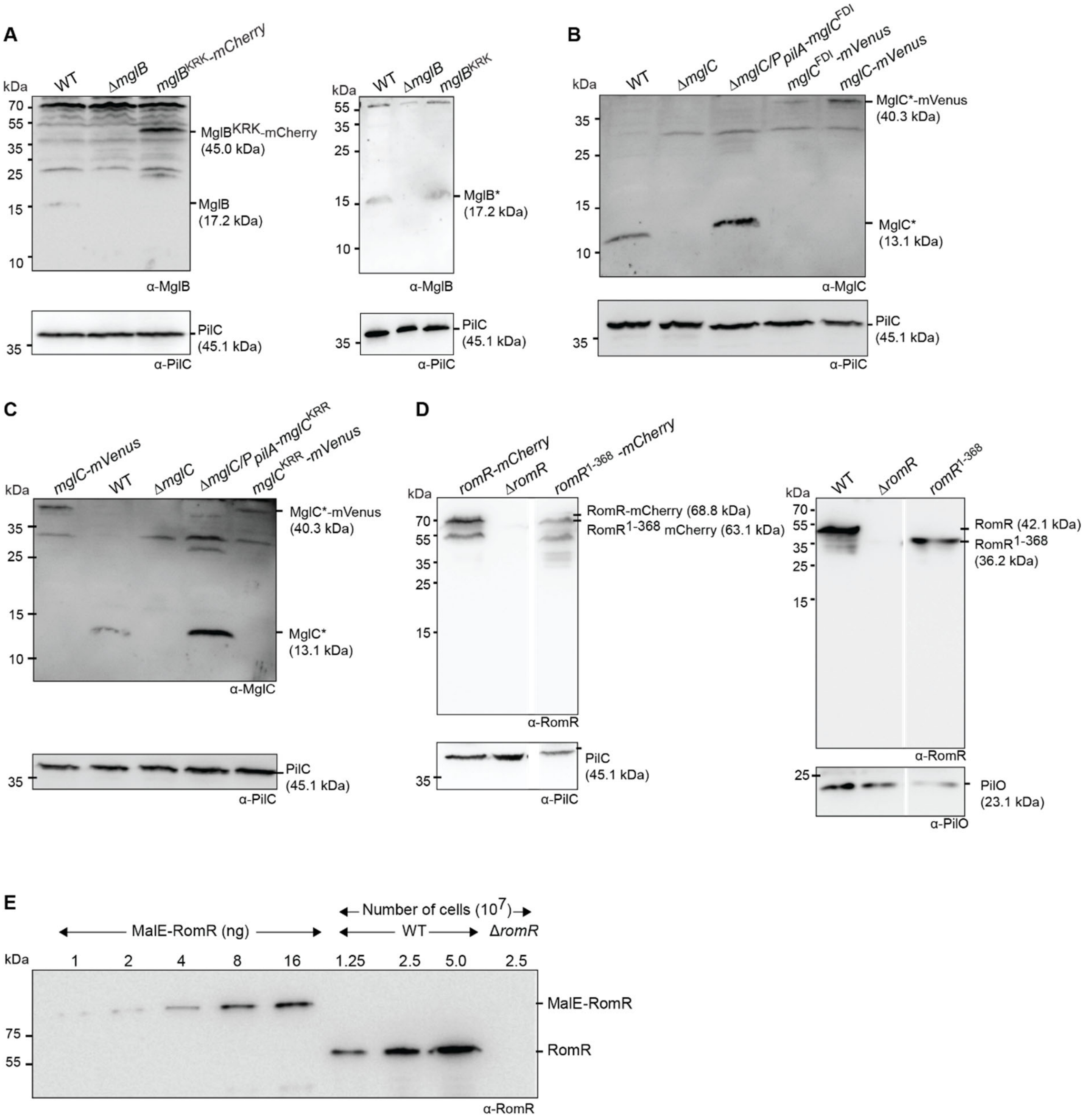


Sup. Fig. 5. Analysis of the accumulation of MglB, MglC and RomR variants

A-D. Immunoblot analysis of accumulation of MglB and MglB-mCherry variants (A), MglC and MglC-mVenus variants (B-C), and RomR and RomR-mCherry variants (D). PilC or PilO were used as a loading controls. Immunoblot analysis was done as in Fig. S1B. All mutant alleles were expressed from the relevant native site except for those labelled P*pilA*, which were expressed from the *pilA* promoter on a plasmid integrated in a single copy at the Mx8 *attB* site.

E. Quantification of RomR molecules per cell by immunoblotting. Cell extracts from the indicated number of WT cells or Δ*romR* cells were probed with specific α-RomR antibodies in parallel with different amounts of purified MalE-RomR. Molecule number per cell was calculated from the intensity of the band in WT lysates compared to a standard curve prepared from the dilution series of known amounts of MalE-RomR on the same blot. Concentrations were calculated based on three independent immunoblots. Shown is a representative blot.

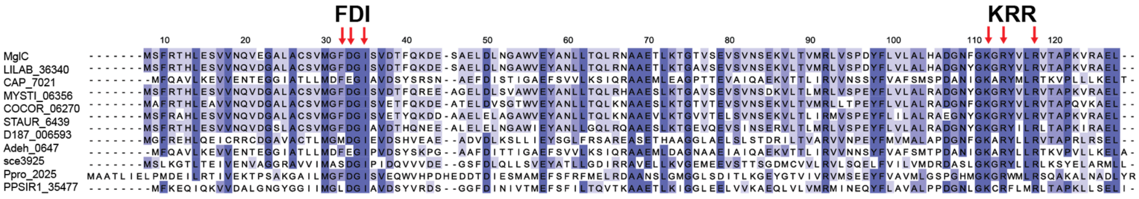

Sup. Fig. 6. Alignment of MglC homologs

Identity is indicated in shades of blue. The FDI and KRR residues are indicated with red arrows.

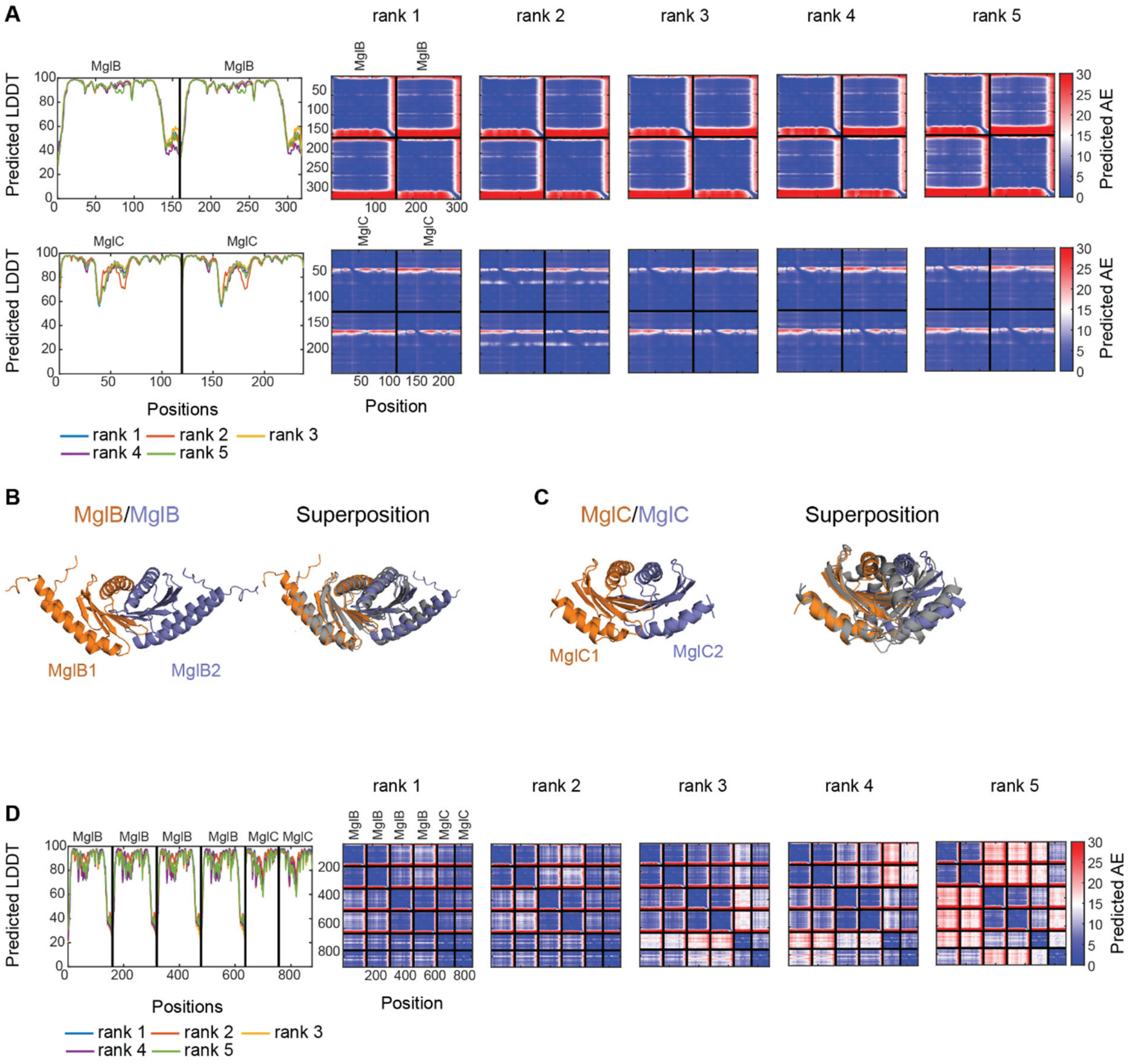


<u>Sup. Fig. 7. AlphaFold-Multimer models of dimeric MglB, dimeric MglC and the $(MglB_2)_2$:$MglC_2$ complex.</u>

A. pLDDT and pAE plots of five AlphaFold-Multimer models of dimeric MglB and dimeric MglC.

B. Superposition of the rank 1 AlphaFold-Multimer model of dimeric MglB (orange and blue) on the solved structure of MglB (pdb ID: 6IZW[3]) (grey).

C. Superposition of the rank 1 AlphaFold-Multimer model of dimeric MglC (orange and blue) on the solved structure of MglC (pdb ID: 7CY1[4] (grey).

D. pLDDT and pAE plots of five AlphaFold-Multimer models of the $MglC_2$:$(MglB_2)_2$ complex. The Rank 1 model is the one used in Fig. 6A and Fig. 7C.

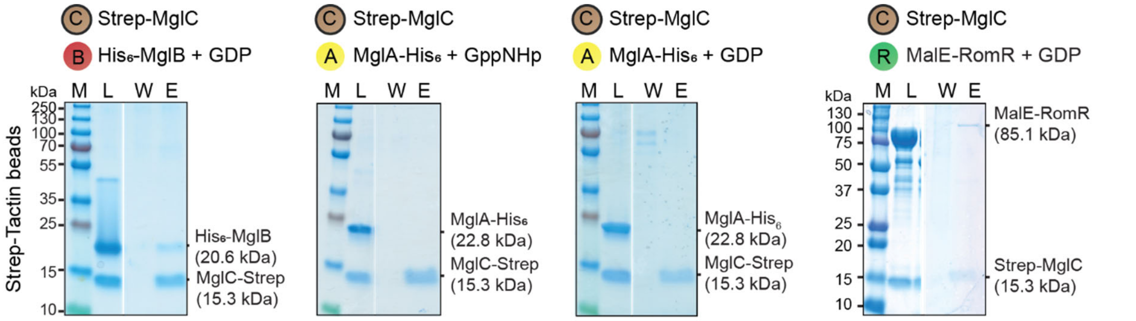


Sup. Fig. 8. *In vitro* pull-down experiments with MglA-His$_6$, His$_6$-MglB, Strep-MglC and MalE-RomR

Control experiments to the experiments shown in Fig. 7AB. Proteins were mixed as indicated at final concentrations of 10µM and applied to the indicated matrix. The matrix was washed and bound proteins eluted. The bait protein is indicated by the black circle. In experiments with MglA-His$_6$, the protein was preloaded with the indicated nucleotide and all buffers contained 40µM of the relevant nucleotide. Equivalent volumes of the load (L), last wash (W) and elution fraction (E) were separated on the same SDS-PAGE gel and stained with Coomassie Brilliant Blue. Gap between lanes indicates lanes deleted for presentation purposes. Calculated molecular masses of the indicated proteins are indicated on the right. Molecular weight markers (M) are indicated on the left.

**Supplementary Table 1.** *M. xanthus* strains used in this work

| Strains | Genotype | Source or Reference |
|---|---|---|
| DK1622 | WT | [5] |
| DK10410 | Δ*pilA* | [6] |
| SA5293 | Δ*aglQ* | [7] |
| SA7300 | Δ*mglC* | [8] |
| SA4420 | Δ*mglA* | [9] |
| SA3387 | Δ*mglB* | [10] |
| SA3300 | Δ*romR* | [11] |
| SA7301 | Δ*mglA* Δ*mglC* | [8] |
| SA7302 | Δ*mglB* Δ*mglC* | [8] |
| SA7304 | Δ*romR* Δ*mglC* | [8] |
| SA8802 | Δ*frzE* | [12] |
| SA7305 | Δ*mglC* $P_{pilA}$-*mglC* | [8] |
| SA8185 | *mglA-mVenus* | [12] |
| SA8130 | *mglA-mVenus* Δ*mglC* | This work |
| SA6963 | *mglB-mCherry* | [11] |
| SA8155 | *mglB-mCherry* Δ*mglC* | This work |
| SA7507 | *romR-mCherry* | [12] |
| SA8129 | *romR-mCherry* Δ*mglC* | This work |
| SA10466 | *mglC-mVenus* | This work |
| SA10627 | *mglC-mVenus* Δ*mglA* | This work |
| SA10535 | *mglC-mVenus* Δ*mglB* | This work |
| SA10467 | *mglC-mVenus* Δ*romR* | This work |
| SA10627 | *mglC-mVenus* Δ*mglA* | This work |
| SA10891 | *mglC-mVenus* Δ*mglA* Δ*mglB* | This work |
| SA10925 | *mglC-mVenus* Δ*mglA* Δ*romR* | This work |
| SA10954 | *mglC-mVenus* Δ*mglA* Δ*mglB* Δ*romR* | This work |
| SA3971 | *mglB-mCherry* Δ*mglA* | [11] |
| SA10971 | *mglB-mCherry* Δ*mglA* Δ*mglC* | This work |
| SA10776 | *mglB-mCherry* Δ*mglA* Δ*romR* | [2] |
| SA12836 | *mglB-mCherry* Δ*mglA* Δ*mglC* Δ*romR* | This work |
| SA7579 | *romR-mCherry* Δ*mglA* | [12] |
| SA10788 | *romR-mCherry* Δ*mglA* Δ*mglB* | [2] |
| SA10898 | *romR-mCherry* Δ*mglA* Δ*mglC* | This work |
| SA11027 | *romR-mCherry* Δ*mglA* Δ*mglB* Δ*mglC* | This work |
| SA10807 | *pVan-romR-mCherry* Δ*mglA* Δ*mglB* Δ*romR* Δ*frzE* Δ*aglQ* | [2] |
| SA12814 | *pVan-romR-mCherry* Δ*mglA* Δ*mglB* Δ*mglC* Δ*romR* Δ*frzE* Δ*aglQ* | This work |
| SA10769 | *pVan-romR-mCherry* Δ*mglA* Δ*romR* Δ*frzE* Δ*aglQ* | [2] |
| SA12899 | *pVan-romR-mCherry* Δ*mglA* Δ*mglC* Δ*romR* Δ*frzE* Δ*aglQ* | This work |
| SA12895 | $mglB^{K14A\ R115A\ K120A}$-*mCherry* | This work |
| SA12885 | $mglB^{K14A\ R115A\ K120A}$-*mCherry* Δ*mglA* | This work |
| SA12896 | $mglB^{K14A\ R115A\ K120A}$-*mCherry* Δ*mglC* | This work |
| SA12894 | *mglA-mVenus* $mglB^{K14A\ R115A\ K120A}$ | This work |
| SA12897 | *mglC-mVenus* $mglB^{K14A\ R115A\ K120A}$ | This work |
| SA12898 | *romR-mCherry* $mglB^{K14A\ R115A\ K120A}$ | This work |
| SA11080 | $mglC^{F25A\ D26A\ I28A}$-*mVenus* | This work |
| SA12893 | *mglA-mVenus* Δ*mglC* $P_{pilA}$-$mglC^{F25A\ D26A\ I28A}$ | This work |
| SA12869 | *mglB-mCherry* Δ*mglC* $P_{pilA}$-$mglC^{F25A\ D26A\ I28A}$ | This work |
| SA12817 | *romR-mCherry* Δ*mglC* $P_{pilA}$-$mglC^{F25A\ D26A\ I28A}$ | This work |
| SA12892 | $mglC^{K104A\ R106A\ R110A}$-*mVenus* | This work |
| SA12887 | $mglC^{K104A\ R106A\ R110A}$-*mVenus* Δ*mglA* | This work |
| SA12890 | $mglC^{K104A\ R106A\ R110A}$-*mVenus* Δ*mglB* | This work |
| SA12899 | $mglC^{K104A\ R106A\ R110A}$-*mVenus* Δ*romR* | This work |

| | | |
|---|---|---|
| SA12886 | *mglC*$^{K104A\ R106A\ R110A}$-*mVenus* Δ*mglA* Δ*mglB* | This work |
| SA12889 | *mglA-mVenus* Δ*mglC* $P_{pilA}$-*mglC*$^{K104A\ R106A\ R110A}$ | This work |
| SA12891 | *mglB-mCherry* Δ*mglC* $P_{pilA}$-*mglC*$^{K104A\ R106A\ R110A}$ | This work |
| SA12888 | *romR-mCherry* Δ*mglC* $P_{pilA}$-*mglC*$^{K104A\ R106A\ R110A}$ | This work |
| SA10955 | *romR*$^{1-368}$-*mCherry* | This work |
| SA10921 | *romR*$^{1-368}$, *mglA-mVenus* | This work |
| SA10966 | *romR*$^{1-368}$, *mglB-mCherry* | This work |
| SA11057 | *romR*$^{1-368}$, *mglC-mVenus* | This work |

**Supplementary Table 2.** Plasmids used in this work

| Plasmids | | Source or reference |
|---|---|---|
| pBJ114 | In-frame deletion vector, $Km^R$ | [13] |
| pSW105 | Complementation from *attB* with $P_{pilA}$, $Km^R$ | [14] |
| pAM5 | $P_{pilA}$ *mglC* Mx8 *attP*, $Km^R$ | [8] |
| pLC66 | pBJ114, construct for *mglC* replacement by *mglC-mVenus* at native site, $Km^R$ | This work |
| pLC20 | pBJ114, construct for *mglA* replacement by *mglA-mVenus* at native site, $Km^R$ | [12] |
| pDK145 | pBJ114, construct for *mglB* replacement by *mglB-mCherry* at native site, $Km^R$ | [11] |
| pLC32 | pBJ114, construct for *romR* replacement by *romR-mCherry* at native site, $Km^R$ | [12] |
| pAM1 | pBJ114 Δ*mglC,* construct for generation of in-frame deletion of *mglC*, $Km^R$ | [8] |
| pSL16 | pBJ114 Δ*mglA,* construct for generation of in-frame deletion of *mglA*, $Km^R$ | [9] |
| pES2 | pBJ114 Δ*mglB* construct for generation of in-frame deletion of *mglB,* $Km^R$ | [10] |
| pSL37 | pBJ114 Δ*romR*, construct for generation of in-frame deletion of *romR,* $Km^R$ | [11] |
| pBJΔ*aglQ* | pBJ114 Δ*aglQ*; construct for generation of in-frame deletion of *aglQ,* $Km^R$ | [15] |
| pLC155 | pBJ114, construct for generation of in-frame deletion of Δ369-420 of *romR,* $Km^R$ | This work |
| pLC154 | pBJ114, construct for generation of in-frame deletion of Δ369-420 from *romR-mCherry,* $Km^R$ | This work |
| pLC301 | pBJ114, construct for *mglB* replacement by $mglB^{K14A\ R115A\ K120A}$ at native site, $Km^R$ | This work |
| pLC280 | pBJ114, construct for *mglB* replacement by $mglB^{K14A\ R115A\ K120A}$-*mCherry* at native site, $Km^R$ | This work |
| pLC250 | $P_{pilA}$ $mglC^{F25A\ D26A\ I28A}$ Mx8 *attP*, $Km^R$ | This work |
| pLC260 | $P_{pilA}$ $mglC^{K104\ R106\ R110}$ Mx8 *attP*, $Km^R$ | This work |
| pLC295 | pBJ114, construct for *mglC* replacement by $mglC^{F25A\ D26A\ I28A}$-*mVenus* at native site, $Km^R$ | This work |
| pLC263 | pBJ114, construct for *mglC* replacement by $mglC^{K104A\ R106A\ R110A}$-*mVenus* at native site, $Km^R$ | This work |
| pLC1 | $P_{van}$ *romR-mCherry*, vanillate-dependent expression of *romR-mCherry* from *mxan18-19* locus, $Tc^R$ | [2] |
| pASKIBA15+ | Vector for overexpression of Strep-tagged proteins; $Km^R$ | IBA Life-sciences GmbH |
| pLC242 | Overexpression Strep-MglC, $Amp^R$ | This work |
| pDK28 | Overexpression MalE-RomR, $Amp^R$ | [11] |
| pTM1 | Overexpression MglA-$His_6$, $Km^R$ | [16] |
| pTM2 | Overexpression $His_6$-MglB, $Km^R$ | [16] |
| pLC283 | Overexpression $His_6$-$MglB^{K14A\ R115A\ K120A}$, $Km^R$ | This work |
| pLC300 | Overexpression MalE-$RomR^{1-368}$, $Amp^R$ | This work |
| pLC310 | Overexpression MalE-MglC, $Amp^R$ | This work |

**Supplementary Table 3**. Primers used in this work

| **Primers (5'-3')** | |
|---|---|
| MglC_E | TTGGTGAAGCCCCCGTAACA |
| MglC_F | CTTGCCATTGTAGAAGAGGA |
| MglC_G | AGC GTG ATG GGC TTT GAC GGC ATC |
| MglC_H_2 | TAC CAT CCG CGT GAA GGG CGA GCA CCA |
| MglC_FW | ATCG AAG CTT AGG CCA CGT ACC CCG TCA |
| MglC_Link_RV | CTC GCC CTT GCT CAC CAT GAG CTC GGC GCG CAC CTT |
| MglCDS800_FW | ATCG TCT AGA TCG GAT GCC CGG CCG |
| MglCDS800_RV | ATCG GAA TTC CGC CTG GGC CCG GGT |
| RomR_FW_HindIII | ATCG AAG CTT CGC CGG GGG CCC GTC |
| RomR_Cterm_B | CCA GGC GCC TCA AGG CGC ACG GGC GCT CGC CGG |
| RomR_Cterm_C | GCG CCT TGA GGC GCC TGG CGC CGT AAC CTC |
| RomR_dsRV_EcoRI | ATCG GAA TTC ATC AGG TCC TGG TAG CGC TCG TC |
| R_Cterm_B_linkerlessgc | TGA TCC ACC GCC TCC AGG CGC ACG GGC GCT CGC CGG |
| Venus_FW_LessGC | GGA GGC GGT GGA TCA ATG GTG AGC AAG GGC GAG |
| GFP RV EcoRI | ATCG GAA TTC TTA CTT GTA CAG CTC GTC CAT GCC G |
| MglC_FA/DA/IA_UP | TGC AGC GTG ATG GGC GCT GCC GGC GCC TCC |
| MglC_FA/DA/IA_DS | GAA GGT GTC GAC GGA GGC GCC GGC AGC GCC |
| MglC_KA/RA/RA_UP | TTC GGC GCG GGC GCC TAC GTG CTG GCC GTC ACT |
| MglC_KA/RA/RA_DS | AGT GAC GGC CAG CAC GTA GGC GCC CGC GCC GAA |
| MglAB_natprom_fw | ATCG AAG CTT CGTCGTTCGGCGTGAAGCCCT |
| mglb_k14_rv | AAA CGG CGT TGA TCG CGG TGA ACT CCT |
| mglb_k14_fw | AGG AGT TCA CCG CGA TCA ACG CCG TTT |
| mglb_r105k120_rv | GTC GCT GGC CGC CTT GAT GCG AAG GGC GAC GAG GC |
| mglb_r105k120_fw | GGC CTC GTC GCC CTT CGC ATC AAG GCG GCC AGC GA |
| MglA_RV_EcoRI | ATC GGA ATT CTC AAC CAC CCT TCT TGA |
| MglC_fw_xbaI | ATC TCT AGA ATG TCC TTC CGC ACG CAC CTC |
| MglC_rv_HindIII | ATC AAG CTT CTA GAG CTC GGC GCG CAC CTT |
| MglC_fw_EcoRI | ATC GAA TTC ATG TCC TTC CGC ACG CAC CTC GAG |
| romr_fw_EcoRI | ATCG GAA TTC ATG CCC AAG AAT CTG CTG GTC GC |
| romr_368_rv_hindIII | ATC AAG CTT AGG CGC ACG GGC GCT CGC CGG |

**Supplementary Table 4.** Quantification of polar fluorescence signals

| Strain | Mean fraction of fluorescence at pole 1 | Mean fraction of fluorescence at pole 2 | Mean fraction of total polar fluorescence | Fraction of cells w/two clusters | Fraction of cells w/one cluster |
|---|---|---|---|---|---|
| *mglC-mVenus* | 0.35 | 0.14 | 0.49 | 0.99 | 0.01 |
| *mglC-mVenus ΔmglA* | 0.57 | 0.05 | 0.62 | 0.79 | 0.20 |
| *mglC-mVenus ΔmglB* | 0.13 | 0.07 | 0.20 | 1.00 | 0.00 |
| *mglC-mVenus ΔromR* | 0.01 | 0.00 | 0.02 | 0.19 | 0.68 |
| *mglA-mVenus* | 0.01 | 0.00 | 0.02 | 0.55 | 0.37 |
| *mglA-mVenus ΔmglC* | 0.00 | 0.00 | 0.00 | 0.09 | 0.38 |
| *mglB-mCherry* | 0.07 | 0.03 | 0.10 | 0.90 | 0.10 |
| *mglB-mCherry ΔmglC* | 0.02 | 0.00 | 0.02 | 0.36 | 0.55 |
| *romR-mCherry* | 0.20 | 0.07 | 0.27 | 0.98 | 0.02 |
| *romR-mCherry ΔmglC* | 0.13 | 0.05 | 0.18 | 0.92 | 0.07 |
| *mglC-mVenus ΔmglA ΔmglB* | 0.20 | 0.12 | 0.32 | 0.98 | 0.02 |
| *mglB-mCherry ΔmglA* | 0.22 | 0.05 | 0.28 | 0.77 | 0.22 |
| *mglB-mCherry ΔmglA ΔmglC* | 0.00 | 0.00 | 0.00 | 0.00 | 0.06 |
| *romR-mCHerry ΔmglA* | 0.49 | 0.08 | 0.57 | 0.91 | 0.09 |
| *romR-mCherry ΔmglB* | 0.09 | 0.04 | 0.13 | 0.96 | 0.04 |
| *romR-mCherry ΔmglB ΔmglC* | 0.08 | 0.04 | 0.13 | 0.92 | 0.08 |
| *romR-mCherry ΔmglA ΔmglB* | 0.14 | 0.06 | 0.20 | 0.89 | 0.11 |
| *romR-mCherry ΔmglA ΔmglC* | 0.11 | 0.05 | 0.16 | 0.92 | 0.08 |
| *romR-mCherry ΔmglA ΔmglB ΔmglC* | 0.08 | 0.04 | 0.12 | 0.92 | 0.08 |
| *mglB-mCherry ΔmglC ΔromR* | 0.00 | 0.00 | 0.00 | 0.00 | 0.07 |
| *mglC-mVenus ΔmglA ΔromR* | 0.04 | 0.00 | 0.04 | 0.25 | 0.69 |
| *mglC-mVenus ΔmglB ΔromR* | 0.01 | 0.00 | 0.01 | 0.18 | 0.67 |
| *mglC-mVenus ΔmglA ΔmglB ΔromR* | 0.00 | 0.00 | 0.00 | 0.00 | 0.01 |
| *$mglC^{FDI}$-mVenus* | 0.21 | 0.09 | 0.29 | 0.97 | 0.03 |
| *mglA-mVenus ΔmglC psw105-$mglC^{FDI}$* | 0.00 | 0.00 | 0.01 | 0.08 | 0.47 |
| *mglB-mCherry ΔmglC psw105-$mglC^{FDI}$* | 0.02 | 0.01 | 0.03 | 0.69 | 0.28 |
| *romR-Cherry ΔmglC psw105-$mglC^{FDI}$* | 0.12 | 0.07 | 0.19 | 0.97 | 0.03 |
| *$mglC^{FDI}$-mVenus ΔmglA* | 0.19 | 0.12 | 0.31 | 1.00 | 0.00 |
| *$mglC^{FDI}$-mVenus ΔmglB* | 0.15 | 0.07 | 0.23 | 0.99 | 0.01 |
| *$mglC^{FDI}$-mVenus ΔromR* | 0.00 | 0.00 | 0.00 | 0.00 | 0.01 |
| *romR-Cherry ΔmglC psw105-$mglC^{KRR}$* | 0.13 | 0.04 | 0.17 | 0.93 | 0.07 |
| *mglA-mVenus ΔmglC psw105-$mglC^{KRR}$* | 0.00 | 0.00 | 0.00 | 0.00 | 0.10 |
| *mglB-mCherry ΔmglC psw105-$mglC^{KRR}$* | 0.04 | 0.01 | 0.05 | 0.76 | 0.24 |
| *$mglC^{KRR}$ mVenus* | 0.01 | 0.00 | 0.01 | 0.04 | 0.69 |
| *$mglC^{KRR}$ mVenus ΔromR* | 0.00 | 0.00 | 0.00 | 0.00 | 0.11 |
| *$mglC^{KRR}$ mVenus ΔmglA* | 0.02 | 0.00 | 0.03 | 0.36 | 0.57 |
| *$mglC^{KRR}$ mVenus ΔmglB* | 0.00 | 0.00 | 0.00 | 0.00 | 0.02 |
| *$mglB^{KRK}$-mCherry* | 0.00 | 0.00 | 0.00 | 0.00 | 0.09 |
| *$mglB^{KRK}$-mCherry ΔmglA* | 0.00 | 0.00 | 0.00 | 0.00 | 0.10 |
| *$mglB^{KRK}$-mCherry ΔmglC* | 0.00 | 0.00 | 0.00 | 0.06 | 0.32 |
| *$romR^{1-368}$-Cherry* | 0.00 | 0.00 | 0.00 | 0.00 | 0.23 |
| *mglA-mVenus $romR^{1-368}$* | 0.01 | 0.00 | 0.01 | 0.06 | 0.54 |
| *mglB-mCherry $romR^{1-368}$* | 0.00 | 0.00 | 0.00 | 0.02 | 0.19 |
| *mglC-mVenus $romR^{1-368}$* | 0.01 | 0.00 | 0.01 | 0.19 | 0.49 |
